# Tomato roots sense horizontal/vertical mechanical impedance and divergently modulate root/shoot metabolome

**DOI:** 10.1101/2021.02.01.429093

**Authors:** Alka Kumari, Sapana Nongmaithem, Sameera Devulapalli, Yellamaraju Sreelakshmi, Rameshwar Sharma

**Author notes:** Author for Correspondence, **E-mail** (RS).

## Abstract

Plant roots encounter coarse environs right after emergence from the seeds. Little is known about metabolic changes enabling roots to overcome the soil impedance. Tomato seedlings grown vertically or horizontally, at increasing hardness, exhibited lateral roots proliferation, shorter hypocotyls, and primary roots. In primary root tips, hardness-elicited loss of amyloplasts staining; induced ROS and NO accumulation. The levels of IBA, zeatin, jasmonates, and salicylic acids markedly differed in roots and shoots exposed to increasing hardness. Hardness lowered IAA and elevated ABA levels, while increased ethylene emission was confined to horizontally-impeded seedlings. The trajectories of metabolomic shifts distinctly differed between vertically/horizontally-impeded roots/shoots. In horizontal roots, amino acids were the major affected group, while in vertical roots, sugars were the major group. Commonly affected metabolites in roots and shoots, trehalose, dopamine, caffeoylquinic acid, and suberic acid, hallmarked the signature for hardness. Increasing hardness lowered *SnRK1a* expression in roots/shoots implying regulation of metabolic homeostasis by the SnRK1 signalling module. Our data suggest that though hardness is a common denominator, roots sense the horizontal/vertical orientation and correspondingly modulate metabolite profiles.

**Significance statement:** We show that the tomato roots sense the magnitude of hardness as well as the horizontal and vertical orientation. The hardness divergently modulates the phytohormone and metabolite levels in roots and shoots. The trajectory of the metabolic shift in vertically-grown seedling distinctly differs from horizontally-grown seedlings. ABA and trehalose were the hallmark of hardness stress and may influence metabolic alteration via the SNRK signalling pathway.

## 1 Introduction

The life cycle of higher plants begins with the sprouting of seeds. The sprouting most often happens underneath the soil, and generally, the root is the first organ to emerge. The architecture of the soil surrounding it strongly influences the subsequent growth of the root. The soil being a compact substratum, the proliferating root’s first task is to overcome the soil hardness. Thenceforth, roots continually adapt and overcome mechanical impedance throughout the plant lifecycle. The above adaptation is visually manifested by alteration in root morphology and branching to ramify in the soil. In general, the morphological changes encompass the reduction of the root growth, thickening of the roots, initiation of lateral roots, and enhanced sloughing of border cells of root caps (Potocka and Szymanowska-Pułka, 2018).

At the cellular level, the thickening of the root is mediated by a reduction in the cell size, particularly epidermal cells (Hanbury and Atwell, 2005), and the formation of additional cortical cell layers (Wilson, Robards and Goss, 1977). The root cap being the site of the growth, the mechanical impedance reduces its size (Souty and Rode, 1987). The cellular changes are also accompanied by the thickening of the cell walls that facilitate the pushing of roots through the compact medium (Wilson and Robards, 1978). Underlying these morphological and cellular responses is sensing of the hardness by the roots and regulation of growth.

The knowledge about the molecular basis of mechanical impedance perception by root is incomplete. The current views envisage that plasma membrane proteins tethered to the cell walls may be responsible for sensing mechanical force (Fruleux. Verger and Boudaoud, 2019). One putative candidate is Mid1-complementing activity (MCA) protein, as roots of Arabidopsis loss-of-function *mca1* and *mca2* mutants fails to penetrate hard agar media (Denness et al., 2011). The mutations in receptor-like kinases *Feronia* also elicit growth defects in Arabidopsis root, including reduced competence to penetrate the hard agar (Shih, Miller, Dai, Spalding and Monshausen, 2014). Another candidate for mechanosensing is the *PIEZO* ion channel localized in the columella and lateral root cap cells. The knockout mutants of the *PIEZO* gene diminished root tips’ capacity to enter hard agar layers (Mousavi et al., 2020).

The root construes the physical impedance to growth as a stress signal. Consistent with this, on the imposition of impedance, one of the earliest responses is enhancing ethylene emission by the plant (Pandey et al., 2020; Sarquis. Jordan and Morgan, 1991). Likewise, the application of exogenous ethylene phenocopies the root morphology akin to the impeded root (Moss, Hall and Jackson, 1988). Conversely, the ethylene signalling inhibitors partially reverse the morphological changes induced by hardness imposition (Sarquis et al., 1991). The ethylene action on root penetration also requires auxin coaction, as loss of root penetration by inhibition of ethylene action can be reversed by auxin (Santisree et al., 2011). The reduction of the root growth in compact soils also affects the shoot growth, indicating coordination between plants’ overall growth responses. The above decline in shoot growth is attributed to the accumulation of ABA in roots and its translocation to shoot, stimulating ethylene emission (Roberts, Hussain, Taylor and Black, 2002; Tracy, Black, Roberts, Dodd and Mooney, 2015).

Unlike obstacle avoidance, wherein roots bend and grow away from obstacles (Lee, Kim, Park, Cho and Jeon, 2020), in compact soils, roots overcome hardness using multiple processes. One such process is enhanced mucilage exudation from the root tips on increased mechanical impedance (Okamoto and Yano, 2017). This entails that hardness elicits the metabolomic alterations in the root, manifested by enhanced mucilage secretion. The metabolome shift is also indicated by enhanced oxygen consumption by mechanically impeded roots, signifying higher respiration rates (Schumacher and Smucker, 1981). Though auxin/ethylene modulates the root penetration response (Santisree et al., 2011), the root metabolome is seemingly not directly linked with auxin/ethylene-induced transcriptome changes (Hildreth, Foley, Muday, Helm and Winkel, 2020). The metabolic activity and flux were instead modulated at the post-translational level by altering enzyme activities and/or transport networks (Hildreth et al., 2020).

Since it is difficult to study the impact of soil compactness on root morphology, the studies in Arabidopsis used seedlings grown horizontally or vertically on agar covered with the dialysis membrane (Okamoto et al., 2008). The above study indicated a role for ethylene signalling in hardness-induced growth inhibition and thickening of roots. However, the dialysis membrane usage confers a constant magnitude of the hardness, while the roots encounter a varying degree of hardness in the soil. To overcome this limitation, we examined the impact of substratum hardness by directly growing tomato seedlings on agarose with varying compactness degrees. We show that hardness impacts hormonal and metabolomic profiles of both root and shoots in a complex fashion. We also show that responses of horizontally and vertically impeded seedlings are considerably different. Our study indicates that the hardness induces extensive metabolic reprogramming, which involves the accumulation of signature metabolites like trehalose, dopamine, and suberin. The above reprogramming is likely modulated by Snf1-related protein kinase1 (*SnRK1*), whose expression is also impacted by the hardness.

## 2 Material and Methods

### 2.1 Plant growth condition

Seeds of tomato *(Solanum lycopersicum)* cultivar Ailsa Craig were surface sterilized with sodium hypochlorite (4% (v/v), 10min). After washing, seeds were sown on moist filter papers and transferred to darkness at ambient temperature for 48h. The sprouted seeds were then sown on agarose (SeaKem LE Agarose, catalog No.50004) (0.2%, 0.6%, 0.8%, 1% and 1.2% all w/v in distilled water) plates (24.0 *l* × 24.0 *w* ×1.2 *h* cm) oriented vertically or trays (48.0 *l* × 36.0 *w* ×9 *h* cm) oriented horizontally (Figure S1A,B). The plates/tray were sealed with parafilm to retain the moisture. The seedlings were grown under 16h light (100 μmole/m^2^/sec) / 8h darkness cycles for 5-days. At the end of 5-day, the seedlings were excised at root/shoot junction, and respective tissue was collected. Collected tissue was snap-frozen, homogenized in liquid nitrogen, and stored at −80°C till further use. The hardness levels of agarose were determined by Durofel DFT 100 (Figure S1C).

### 2.2 Phenotyping and imaging

The root (including lateral roots) and shoot lengths were measured using a ruler. For morphological measurements, ~40 seedlings were observed per replicate, and five independent biological replicates were used. All microscope images were recorded using an Olympus BX-45 fluorescent microscope. For root tip imaging, approximately 1 cm of root tips were used. For DAPI (4’, 6-diamidino-2-phenylindole) staining, excised root tips were incubated in 300 nM solution for 2-4 min in darkness, followed by washing with PBS buffer and imaging using 358nm excitation and 461nm emission wavelength. The NO levels in root tips were monitored by DAF-2 DA (4,5-Diaminofluorescein diacetate) fluorescence as described in Negi, Santisree, Kharshiing and Sharma (2010). For ROS detection, the Pelagio-Flores, Ruiz-Herrera and López-Bucio (2016) procedure was followed. The *IAA2::GUS* staining of root tips/shoots/whole seedlings was performed as described in Santisree et al. (2011). The amyloplasts in root tips were stained as described by Benjamins, Quint, Weijers, Hooykaas and Offringa (2001). For all microscope imaging experiments, ~20 seedlings were examined per biological replicate, and 3-5 independent biological replicates were used.

### 2.3 Ethylene emission

To measure ethylene emission, seedlings were grown in air-tight boxes. Ethylene released from seedlings was adsorbed on 0.25M of mercuric perchlorate soaked cotton. Adsorbed ethylene was released by adding four volumes of 4M NaCl relative to mercuric perchlorate. The ethylene emission was monitored by gas-chromatography as described earlier (Santisree et al., 2011). The total ethylene emission was normalized to the seedlings’ weight and the incubation period with mercuric perchlorate. For ethylene emission, ~20 seedlings were examined per biological replicate, and three independent biological replicates were used.

### 2.4 qRT-PCR

Total RNA was extracted from the roots (including lateral roots) and shoots (hypocotyl and cotyledons) using TRI Reagent (Sigma, USA) using the manufacturer’s protocol. The cDNA synthesis and qRT-PCR was carried out described in Kumari et al. (2017). The list of genes and primer sequences are provided in Table S1. For qRT-PCR, roots/shoots from ~20 seedlings were harvested per biological replicate, and three independent biological replicates were used.

### 2.5 Profiling of phytohormones and metabolites

The endogenous phytohormone levels were determined using Orbitrap Exactive-plus LC-MS following the protocol described earlier (Kumari et al., 2017; Pan, Welti and Wang, 2010). Primary metabolite profiling by GC-MS was carried out by a method modified from Roessner, Wagner, Kopka, Trethewey and Willmitzer (2000) described in Kumari et al. (2017). For the identification of primary metabolites, the analysis was performed as mentioned in Kumari et al. (2017). Table S2 and Table S3 lists the metabolites detected in root and shoot, respectively. Metabolic networks were constructed as described in Kumari et al. (2017). Correlation coefficient (PCC) value ± ≤ 0.95 was used to create networks. For the above experiments, roots/shoots from ~20 seedlings were harvested per biological replicate, and five independent biological replicates were used.

### 2.6 Statistical Analysis

All Statistical Analysis On Microsoft-Excel (http://prime.psc.riken.jp-/MetabolomicsSoftware/StatisticalAnalysisOn-MicrosoftExcel/) was used to obtain significant differences between data sets. Statistically significant differences between treatments were determined by one-way ANOVA using the Student-Newman-Keuls method with P≤ 0.005 and P≤ 0.001.

## 3 Results

### 3.1 Vertical vs. horizontal impedance differentially affects root morphology

To monitor the effect of increasing hardness on root growth, we selected 0.2% agarose (A) media as control. At 0.2%A, agarose forms a very soft gel and imposes minimal hardness stress on the roots. The tomato seeds rest on the surface of 0.2%A (Figure S1B), but roots penetrate and proliferate in the gel. For all experiments, seedlings grown on 0.2%A served as control. We selected ≥0.6%A for hardness experiments, as ≥0.6%A gel was enough firm to hold itself, without slipping, in vertically oriented plates. Also, tomato roots do not penetrate in agarose in horizontally oriented plates. Tomato seedlings grown on ?0.6%A showed typical inhibition of primary root elongation with increasing hardness (Figure 1A-a, Figure S1D). Compared to *v*ertically-grown *s*eedlings (VS), primary root elongation was starkly affected in *h*orizontally-grown *s*eedlings (HS) (Figure 1A). In both HS and VS, the primary root length was reduced to ≥ 50% at 1.2%A. Above hardness-imposed responses were similar to those elicited by soil compaction in tomato seedlings (Tracy et al., 2012). Unlike soil, where roots encounter hardness around their circumference, in our study, the root’s exposure to hardness is limited to the side where it touches agarose. Likewise, VS shoots also touch agarose during their growth.

**Figure 1:**
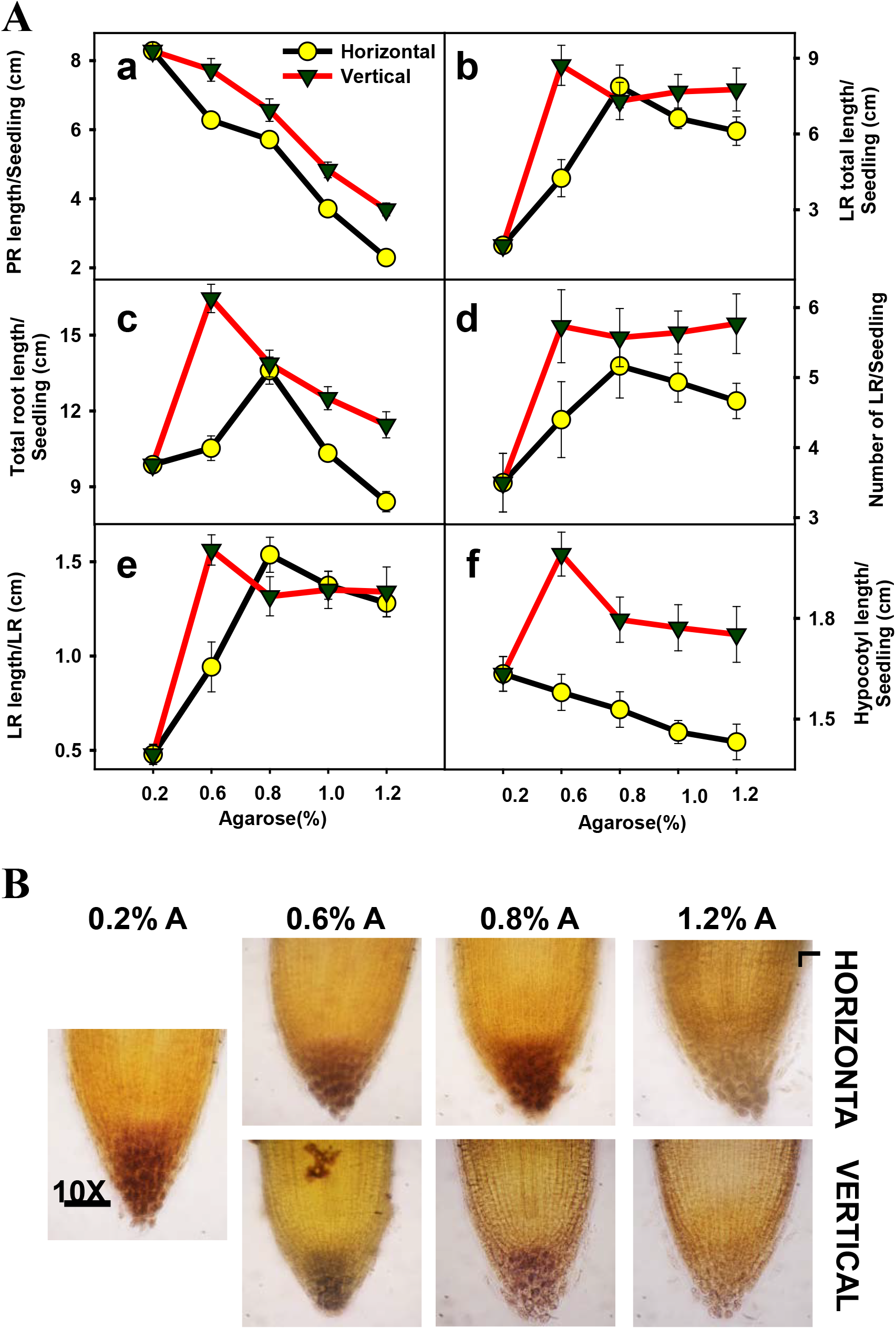
Effect of increasing hardness on root/hypocotyl growth and amyloplasts. **A**, Five-day old seedlings grown in vertically- or horizontally-oriented plates were monitored for morphological changes. **a**, Primary root (PR) length; **b**, Total lateral root (LR) length; **c**, Total root length (Primary + Lateral roots); **d**, Number of lateral roots per seedling; **e**, Mean length of lateral roots per lateral root; **f**, Hypocotyl length. **B.** Progressive loss of amyloplast staining in columella cells of the primary root tip. Note increasing hardness enhanced the sloughing of root cap cells.

In HS, hardness stimulated the initiation and elongation of lateral roots till 0.8%A. Also, in HS, the total length of the lateral roots increased till 0.8%A, then declined (Figure 1A-b,d). Remarkably in HS, for all lateral root-related responses, 0.8%A hardness was an inflection point for reversal. Contrastingly, in VS, beyond 0.8%A inflection point, increasing hardness only marginally affected the number of lateral roots, their total length, and average length (Figure 1A-b,c,d). Nonetheless, more lateral roots with higher length were initiated in VS than HS (Figure 1A-e). The increasing hardness also reduced the hypocotyl length. Again, HS showed higher inhibition of hypocotyl length than VS (Figure 1A-f). Visually, no alteration of cotyledon shape and size was discernible.

The impact of increasing hardness also manifested on the root hairs of HS. With increasing hardness, the initiation of root hairs shifted proximally to the primary root tip. Additionally, the density of root hairs and their length increased in parallel with hardness (Figure S1E). The reduction in root length in HS was associated with a parallel decrease in epidermal cell length. In VS’s root epidermal cells, the cell length decrease was of lesser magnitude (Figure S2). Remarkably, the reduction in root length elicited the formation of additional cell layers in HS (Figure S3A, B). Consistent with this, with increasing hardness the diameter of the HS primary root slightly increased, whereas it remained similar in VS (Figure S3C). Both in HS and VS, increasing hardness also affected amyloplasts staining, which are purported to be associated with gravisensing (Figure 1B). Consistent with increasing hardness, the sloughing-off of columella cells increased in both HS and VS primary roots (Figure 1B). Increased sloughing of root cap cells at higher hardness is consistent with the observations that compacted substratum enhances the sloughing of cap cells (Iijima, Griffiths and Bengough, 2000).

### 3.2 Increasing hardness elevates ROS and NO levels in the primary root tip

It is believed that substratum hardness is perceived primarily by the primary root tip, which is the first organ to emerge from the germinating seeds. Unlike decline in amyloplast staining, the expression of the auxin-responsive reporter *IAA2::GUS* in root tips did not show significant changes with increasing hardness in HS and VS (Figure 2A). The staining of whole seedlings showed localized expression of *IAA2::GUS* in different parts of seedlings. Specifically, higher GUS expression was seen in lateral roots, root/shoot junction, hypocotyl, and part of cotyledons, which seemed to increase with increasing hardness, albeit more in HS (Figure S4A,B). Contrasting to *IAA2::GUS* expression, increasing hardness leads to increased nitric oxide staining in primary root tips (Figure 2B). At lower hardness, the staining was localized in columella cells and its vicinity, and with increasing hardness, it spread distally in the root. A similar increase in the staining was also observed for ROS in root tips, with the region above the quiescent centre staining more deeply than the central cells and lesser staining in epidermal cells (Figure 2C). The deeply stained ROS region seems to shift towards the root tip with increasing hardness in HS and VS.

**Figure 2:**
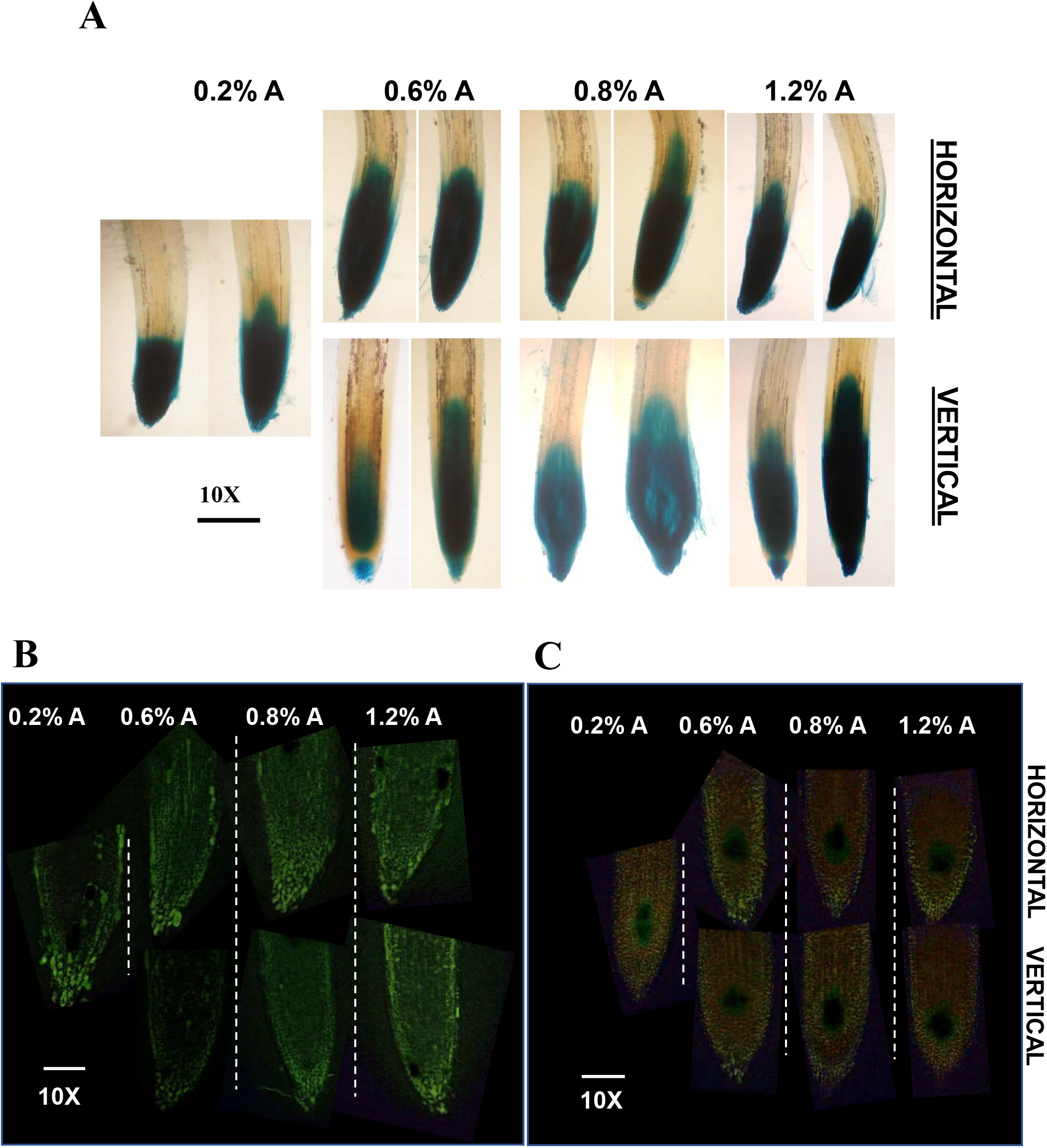
Influence of hardness on *IAA2::GUS,* NO, and ROS levels in primary root tips. **A**, Expression of the auxin reporter *IAA2::GUS* in primary root tips. **B**, DAF-2DA detection of NO levels in primary root tips **C,** DCF-2DA (2’,7’-dichlorofluorescein diacetate) detection of ROS levels in primary root tips. Note increases in NO and ROS level with increasing hardness, unlike *IAA2::GUS,* whose expression is not altered.

### 3.3 Phytohormones are dissimilarly affected in root and shoot

The interrelationship between hardness-induced inhibition of seedling growth and ethylene emission is well-known (Okamoto et al., 2008; Růžička et al., 2007; Swarup et al., 2007). Consistent with this, ethylene emission in HS increased in parallel with a reduction in primary root length. Amazingly, ethylene emission was not affected in VS; despite increasing hardness, the ethylene emission was nearly similar (Figure 3A). Unlike ethylene, whose emission was measured from intact seedlings, the levels of other hormones were estimated from shoots and roots harvested from seedlings.

**Figure 3:**
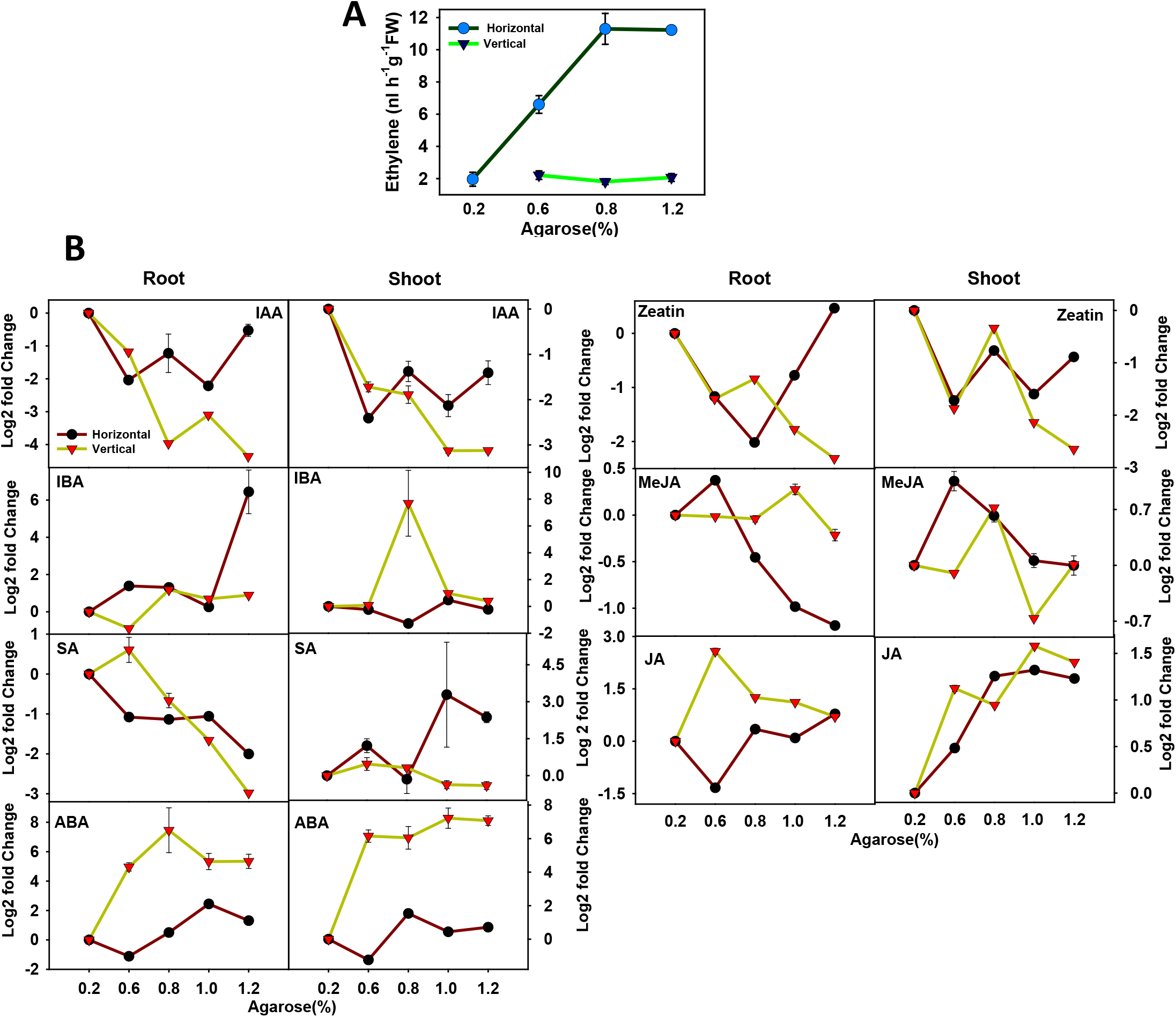
Influence of hardness on phytohormone levels in roots and shoots. The levels of all hormones in roots and shoots (except ethylene) were estimated using Liquid Chromatography-Mass Spectrometry (LC-MS). The ethylene emission from whole seedlings was measured by Gas Chromatography (GC). **A**. Ethylene **B**, Other phytohormones. ABA (Abscisic acid), SA (Salicylic acid), IAA (Indole −3-acetic acid), IBA (Indole-3-butyric acid), JA (Jasmonic acid), MeJA (Methyl Jasmonic acid). The log2 fold changes in respective hormones at different hardness levels (barring ethylene) were determined by calculating the ratio using 0.2%A-grown seedling root/shoot as the denominator. The hormone data, log2 fold, and P values are given in supplementary tables S2.

The hormonal profiles of HS- and VS-roots and shoots were distinctly different (Figure 3B, Table S2). Broadly, most hormones followed different trajectories in shoots than in roots. In VS roots, the IAA level declined with increasing hardness and then remained nearly the same. In HS roots, after a decline at 0.6%A, the IAA level was somewhat stable and rose again at 1.2%A. A similar pattern was seen in shoots. While in HS-shoots, after an initial decline, the IAA level remained relatively stable, it continually declined in VS-shoots. Contrarily in roots, hardness somewhat stimulated IBA levels (barring 0.6%A in VS roots), which was highly exacerbated at 1.2%A in HS roots. Interestingly IBA level was high at 0.8%A in VS shoot. Like IAA, zeatin level first declined and later increased in HS-roots, whereas in VS-roots it was lower but only mildly varied. The zeatin levels were also lower in HS- and VS-shoots.

Among the stress-related hormones, the ABA levels were much higher in VS-roots than HS-roots; it peaked at 0.8%A and 1.0%A, respectively. Similar to roots, VS shoots had higher ABA levels at all hardness levels. Both in HS- and VS-roots, salicylic acid (SA) declined with increasing hardness, which set in earlier in HS than in VS. SA levels were nearly the same in VS-shoots but increased in HS-shoots at higher hardness. The methyl jasmonate (MeJA) accumulation profile also differed in HS- and VS-roots, with a continual decline in the MeJA level in HS-roots with increasing hardness. Contrarily, the MeJA level increased in HS shoots and then declined, while in VS-shoots, its level fluctuated. The jasmonate (JA) levels in HS-roots were nearly the same (barring 0.6%A), while it first attained a high level then decreased in VS roots. Unlike roots, the JA level increased both in VS- and HS-shoots at increasing hardness.

### 3.4 PCA revealed increasing hardness diversely affected metabolite profiles

The GS-MS analysis identified a total of 82 metabolites in roots and shoots (Table S3, S4). The imposition of hardness affected primary metabolite profiles signified by diversely affected PCA profile of root (Figure 4A) and shoot (Figure 4B). In HS-roots, with increasing hardness, the PCA profiles showed a progressive shift in PC2 and a minor shift in PC1 (Figure 4A). Conversely, in VS-roots, increasing hardness led to a progressive shift in PC1 and a minor shift in PC3. Compared to the root, the PCA profiles of shoots at lower hardness showed more overlap, yet these were distinct (Figure 4B). In HS shoots, the PCA profile at 0.6%A was close to control (0.2%A). The PCA profile at 0.8%A showed a shift in PC2. In HS shoots, at higher hardness, the profiles showed a shift both in PC1 and PC3. Conversely, in VS shoots, the PCA profile showed more overlaps and varied minorly along the PC2 axis. Above gradual shifts on diverse planes indicated a significant metabolic divergence between VS and HS roots and shoots.

**Figure 4:**
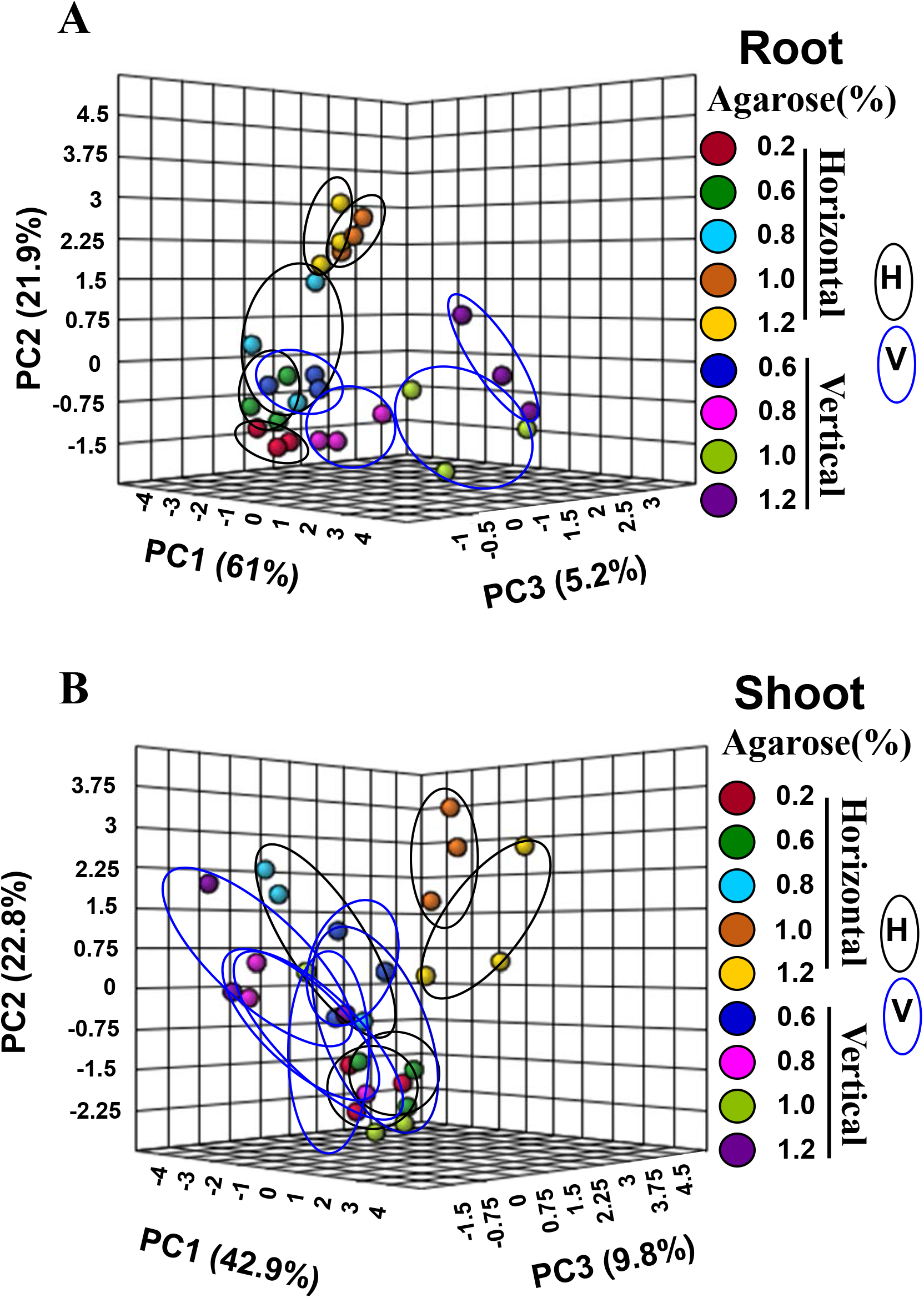
Principle component analysis (PCA) of metabolites of roots (A) and shoots (B). The PCA was constructed using the MetaboAnalyst 3.0. The variance of the PC1, PC2, and PC3 components is given within parentheses. Note PCA profiles of HS- and VS-roots follows different trajectories. Similar to roots, the PCA profile of HS-shoots also shows a wider shift, while that of VS-shoots are in close proximity.

To explore the interactions between metabolites and hormones, the regulatory networks were constructed at high stringency (Pearson’s Correlation Coefficient (PCC) values ± ≤ 0.95) (Figure S5). The Venn diagram revealed that most interactions were unique at a given hardness level. The common interactions steadily declined, with very few interactions being common at all hardness levels (Figure S6A, Table S5A). In general, the networks were in conformity with the PCA that at different hardness metabolites were uniquely modulated, both in root and shoots. Nonetheless, the root networks were markedly modulated by hardness compared to shoots, manifested by a steady increase in negative interactions with increasing hardness (Figure S6B, Table S5B).

### 3.5 Metabolome is divergently affected in VS/HS-roots and shoots

In conformity with distinct PCA profiles, the levels of individual metabolites varied distinctly between roots and shoots (Table S3, S4). The metabolites that were significantly altered (Log2 fold ± ≥1) in two or more hardness levels were only considered changed and are discussed below. In general, the imposition of hardness upregulated (↟) the majority of metabolites in roots and shoots (Figure 5). Conversely, only a few metabolites were significantly downregulated (↓) (HS root-2↓; VS root-2↓; HS shoot-5↓; VS-shoot 4↓). In HS roots, 26 metabolites were upregulated, of which amino acids (12) constituted the majority. In general, at 0.8%A, the upregulation of metabolites in HS root was milder than other hardness levels. Divergently in VS roots, among 28 upregulated metabolites, the majority were sugars (8). Out of 5 upregulated amino acids in VS root, only alanine, oxoproline, and asparagine were common with HS roots. Also, out of 3 TCA cycle intermediates, citric acid and malic acid were common in VS and HS roots. Both HS and VS roots also had common upregulation of different forms of calystegine. Similar to roots, VS shoots (18↟) and HS shoots (16↟) also showed divergence in upregulated metabolites, with few common metabolites. Remarkably in shoots, the TCA cycle intermediates were not upregulated, and only a few amino acids were upregulated (HS 3↟; VS 2↟). Like VS roots, VS shoots also had more upregulated sugars (6↟) than HS (2↟).

**Figure 5:**
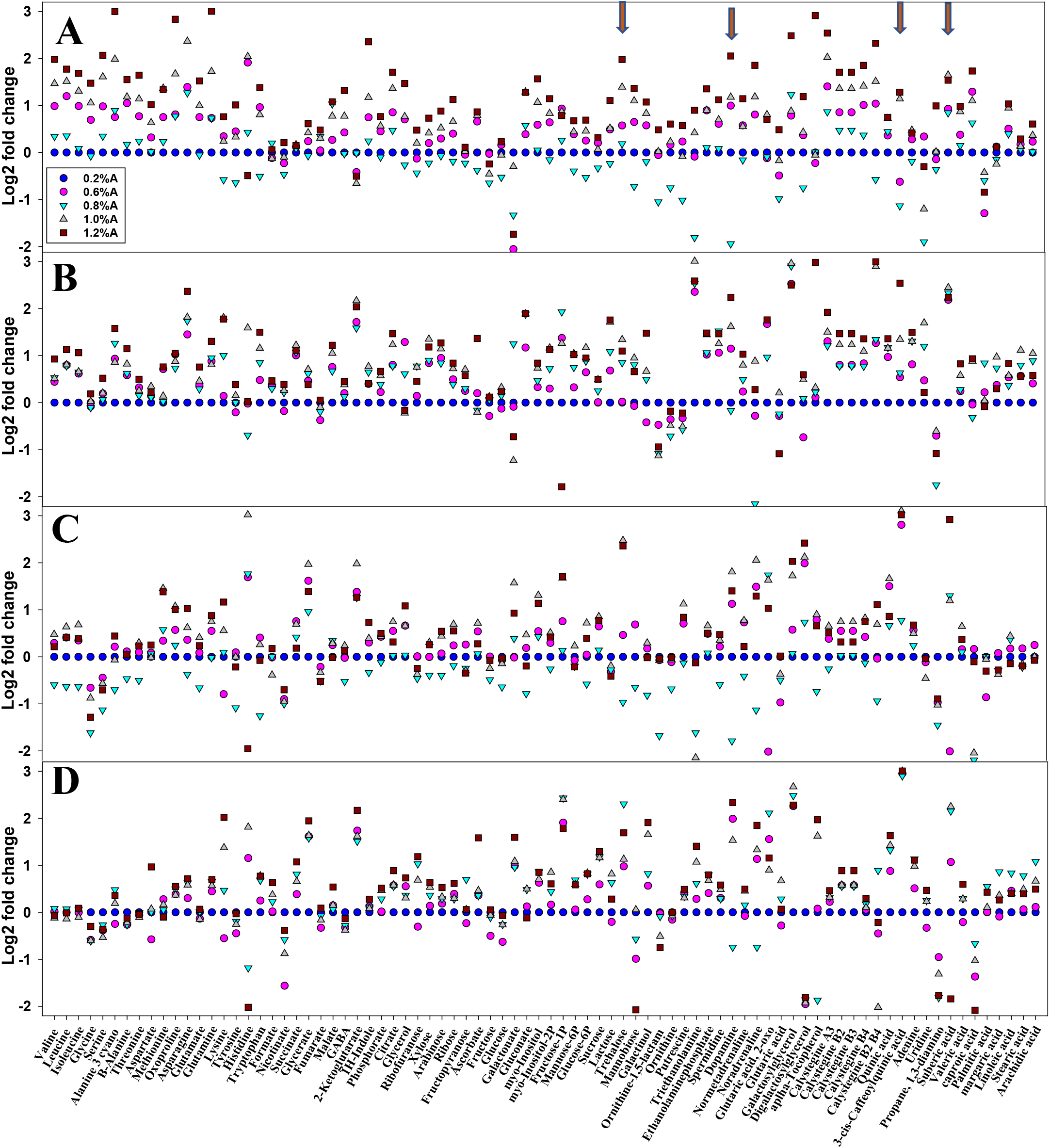
The metabolic shifts in roots and shoots of vertically- or horizontally-oriented seedlings. The log2 fold changes in respective metabolites at different hardness were determined by calculating the ratio using 0.2%A-grown seedling root/shoot as the denominator. **A**, Horizontal roots. **B**, Vertical roots. **C**, Horizontal shoots. **D**, Vertical shoots. The arrows show the four signature metabolites upregulated in roots and shoots of horizontally-or vertically-oriented seedlings. The metabolite data, log2 fold, and P values are given in supplementary tables S3 and S4.

Both in root and shoots, though the levels of several metabolites levels altered, we considered the metabolites upregulated at two or more hardness levels in all four tissues as signature metabolites for hardness. The four metabolites that showed common upregulation in HS/VS roots and shoots were trehalose, dopamine, caffeoylquinic acid, and suberic acid (Figure 5, arrows in A). The metabolites upregulated in HS/VS roots and also in HS shoot were oxoproline and myoinositol. Alike among the upregulated metabolites in HS/VS shoot and common with VS roots were glutonic acid, fructose 1,6 phosphate, 2-ketoglutarate, and that with HS roots was noradrenaline. The metabolite common with both VS root and shoot was triethanolamine (downregulated in HS shoot).

Similarly, methionine-a precursor of ethylene, and histidine were also upregulated in HS roots and shoots. In VS and HS shoots, the level of propane-1-3 diamine (also downregulated in VS-roots) and hexanoic acid (Upregulated in HS-roots) was downregulated. Other differentially regulated metabolites were mannobiose (HS roots↟, shoots↓), galactonic acid (HS roots↓, VS-shoots ↟), and uridine (HS roots ↓, VS roots ↟).

### 3.6 Higher hardness downregulates *SnRK1a* expression

The trigger for the shift in the metabolome in our study was increasing with substratum hardness. Little is known how plant roots sense the hardness, barring that Arabidopsis *mca1* mutant roots fail to penetrate the hard agar medium (Nakagawa et al. 2007). Unexpectedly, the imposition of hardness did not stimulate *MCA1* expression; instead, it lowered its expression in HS-(1.2%A) and VS-roots (1%, 1.2%A). Similarly, HS-shoots, too, showed lower expression (at 1%A). However, VS-shoots showed stimulation of *MCA1* at 1%A (Figure 6. Table S6). Emerging evidences have indicated that *HY5* acts as a coordinating signal between root and shoot. Also, Arabidopsis *hy5* mutants show higher lateral root initiation and loss of wavy response on inclined plates (Oyama, Shimura and Okada, 1997). *HY5* expression declined in roots and shoots at higher hardness, and albeit reduction in VS-shoots was stronger than HS-shoots (Figure 6). The expression of *MYB36,* which plays a role in root growth, declined at the higher hardness in both roots and shoots. The expression of *E2Fa*, which regulates entry into the G-phase of the cell cycle, is lowered in roots and shoots of HS/VS at higher hardness (Figure 6).

**Figure 6:**
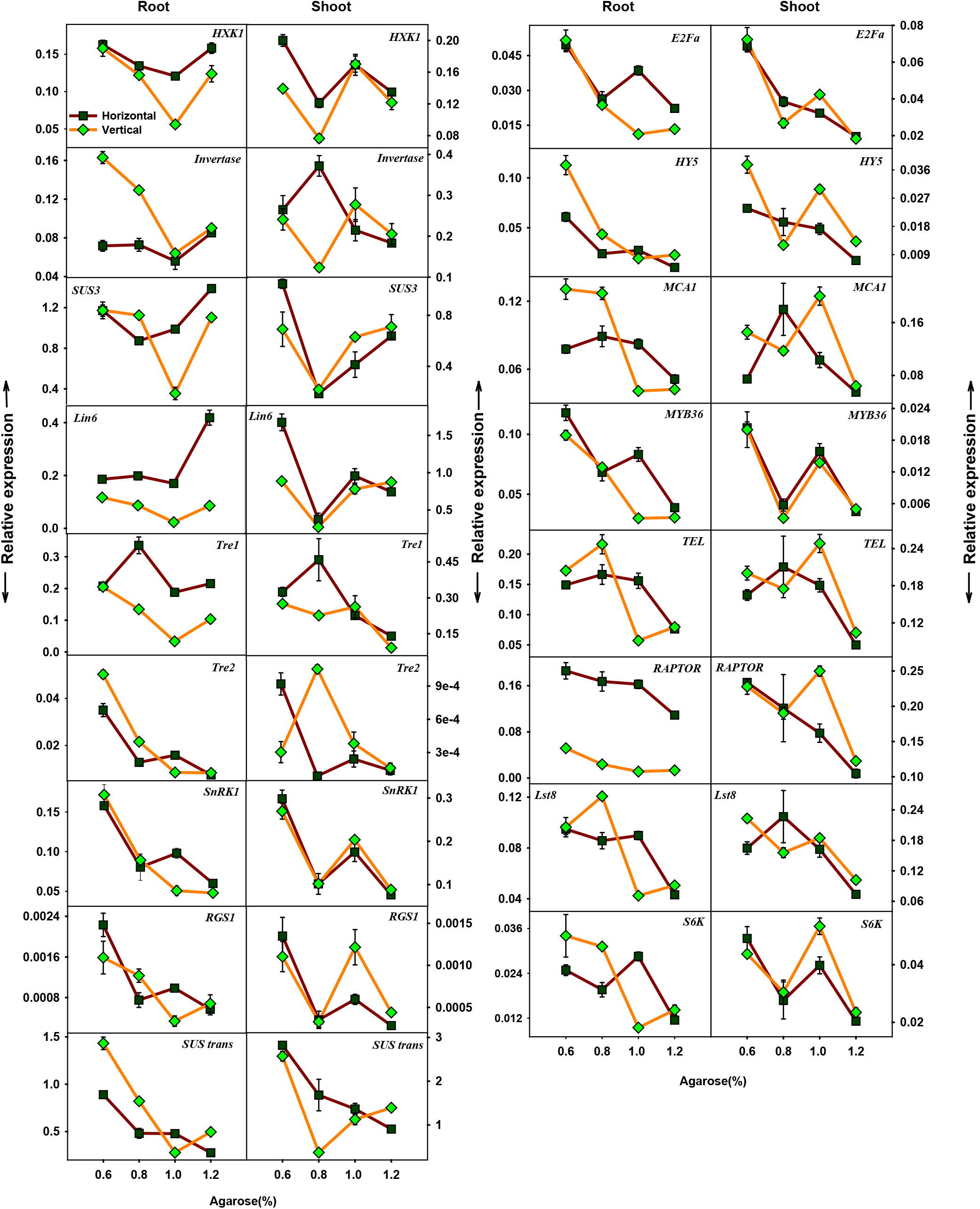
Influence of hardness on the expression of C/N metabolism genes. The expression of genes associated with TOR and SnRK signalling, energy sensing, sugar transport and metabolism, and related transcription factors were examined in root and shoot. *TEL-* target of rapamycin/TOR; Raptor-Regulatory-associated protein of TOR; *Lst8-* Target of rapamycin complex subunit LST8, *S6k-* Ribosomal protein S6 kinase, TOR associated protein; *SnRK1-SNF1*-related protein kinase 1; *HXK1-* Hexokinase1; *Invertase-* Invertase 8; *Lin6-* Invertase; *SUS trans-* Sucrose transporter; *SUS3-* Sucrose synthase; *Tre1-* Trehalose-6-Phosphatase (Shoot specific); *Tre2-* Trehalose-6-Phosphatase (Root specific); *RGS1-* Regulator of G protein signalling; *MCA1*-mid1-complementing activity 1; *E2Fa-* E2Fa transcription factor; *HY5*-Elongated Hypocotyl 5; *Myb36*-Myb domain protein 36.

Metabolite profiling of roots indicated a shift in C/N metabolism, as higher hardness stimulated amino acids and sugar accumulations in HS-roots and VS-roots, respectively. Considering sugar also acts as a signalling molecule, we examined the expression of genes related to sugar signalling, transport, and metabolism viz. *HXK1, RGS1, SUS trans, Lin6, SUS3,* and *invertase.* The *SUS trans* expression was lower at ≥0.8%A in both roots and shoots. While in VS-roots, expression of *invertase* was reduced, it was not affected in HS-roots. In VS/HS-shoots, expression of *invertase* showed the opposite effect at 0.6%A. Expression of *HXK1* was lower at 0.8 and 1.0%A in roots, whereas in shoots, lower expression was shared only at 0.8%A. Higher hardness lowered *SUS3* expression in HS-root (barring 1.2%A) and HS-shoots, but in VS, it was seen only at specific hardness (Roots-1%A; Shoots-0.8%A). Interestingly, *LIN6* was massively upregulated in HS-roots at 1.2%A, while in VS-root (1 and 1.2%A), and HS-shoot (0.8-1.2%A), and VS shoot (0.8%A), its expression was low. The expression of glucose sensor *RGS1*, also a G-protein signalling effector, was lower at the higher hardness in HS and VS roots and shoot (Except VS-root 0.8%A; HS-shoot 1%A) (Figure 6).

Metabolome analysis revealed that hardness stimulated trehalose accumulation. Fortuitously trehalose-6-phosphate (T6P) is also implicated as one of the regulators of metabolic processes. The hardness also affected the expression of the trehalose-6-phosphatase genes (*Tre1-*shoot-specific; *Tre2-* root-specific). Oddly, higher hardness lowered *Tre1* expression in VS-root, also in VS-shoot (0.8, 1.2%A) and HS-shoot (1.2%A). Contrarily, *Tre2* expression declined at the higher hardness in VS- and HS-roots, as well in HS-shoot, while VS-shoots showed enhanced levels at 0.8%A (Figure 6).

Considering that SnRK and TOR signalling influence the metabolomic homeostasis in plants, we examined the relationship between increasing hardness and expression of genes viz., *SnRk1a,* and *TEL, RAPTOR, Lst8,* and *S6K* related to TOR-signalling (Figure 6). Remarkably, increasing hardness lowered the expression of *SnRK1a* in VS as well as HS roots and shoots. Interestingly, both in roots and shoots of HS/VS, the expression of *TEL, RAPTOR, Lst8,* and *S6K* was low at 1.2%A. Also, VS-roots showed lower expression of *TEL, RAPTOR, Lst8,* and *S6K* at 1.0%A. The shoots also showed lower expression of these genes at few hardness levels *(RAPTOR,* HS 1%A, VS 0.8%A; *LST8,* VS 8%A; *S6K,* HS, and VS 0.8%A). An exception was a higher expression of *S6K* in VS-shoots at 1.0%A.

## 4. Discussion

The ramification of roots in the soil involves constant interaction with its ambient environment, as the roots forage the soil for water and mineral nutrients. Parallelly, the roots have to overcome the soil compaction or hardness continually. We examined whether the roots perceive hardness differently when they encounter it in the vertical or horizontal orientation and how the above sensing reprograms hormonal balance and metabolite profiles.

The soil hardness is a well-recognized factor influencing the elongation and proliferation of roots in the soil. The Arabidopsis primary roots grown horizontally on dialysis membrane overlaid on agar displayed reduced growth, cell elongation, and radial thickening. Conversely, roots grown on vertical plates with or without dialysis membrane did not display these alterations. It was assumed that in vertically oriented plates, physical hardness’s imposition did not affect root growth (Okamoto et al., 2008). Contrarily, in tomato, irrespective of the plate’s orientation, hardness inhibited primary root elongation. The parallel reduction in primary root elongation with increasing hardness confirms that the roots sense the substratum hardness, even in the vertical orientation.

The reduced root length is physiologically disadvantageous to growing plants. VS offset this by initiating more and longer lateral roots at all hardness levels. HS also offset it by increased lateral roots initiation and elongation till 0.8%A. The increased root diameter, enhanced root hair number/elongation, and higher reduction in HS’s epidermal cell lengths are akin to morphological changes in roots subjected to mechanical stress (Potocka and Szymanowska-Pułka, 2018). Increasing hardness also impacted hypocotyl elongation, albeit moderately than primary roots. Like primary roots, the reduction of hypocotyl length was more severe in HS than VS, suggesting a causal linkage between roots and shoot growth.

The natural inclination of primary root growth is along the gravitational vector. Since VS-roots can freely grow downwards, these do not encounter gravitational stress. Contrarily, the HS-root, particularly the primary root tip, is constantly under gravitational stress due to its horizontal position. It is believed that sedimentation of amyloplasts localized in columella cells of the root tip in the direction of gravity is critical for gravisensing (Hashiguchi, Tasaka and Morita, 2013). The observed reduction in amyloplasts staining in HS-root tips with increasing hardness can presumably occur either due to gravitational stress imposed by horizontal orientation or by the stress imposed by substratum hardness. Considering that VS-root tips too show a staining reduction, the above reduction seems to be caused by hardness than the gravitational stress. A similar loss of amyloplasts was observed when roots were forced to follow hydrotropism, wherein loss of amyloplasts was attributed to suppression of gravitropism (Takahashi, Yamazaki, Kobayashi, Higashitani and Takahashi, 2003). But in our study, there is no specific reduction in water status; therefore, water stress can be ruled out. It is also reported that auxin stimulates the amyloplast formation in Arabidopsis’ root tips (Zhang et al., 2019). Considering that *IAA2::GUS* staining is nearly uniform, it rules out that the depletion of auxin leads to amyloplast loss.

It is believed that the root tip is the site for sensing the ambient environment in the soil, such as gravity, water, mineral nutrients, and light (Arnaud, Bonnot, Desnos and Nussaume, 2010). Reduced amyloplasts staining indicated that the root tip is either a likely site that also perceives the substratum hardness or reduced staining is merely a stress-induced response. Since VS root tips also show reduced staining, it is unambiguous that the root tips are sensitive to hardness-stress. The higher accumulation of reactive oxygen species (ROS) in cells/organs is considered a hallmark of abiotic stress (Farnese, Menezes-Silva, Gusman and Oliveira 2016). The increased ROS and NO staining of HS and VS root tips are consistent with the assumption that hardness is perceived as a stress signal (Jacobsen. Jervis, Xu, Topping and Lindsey, 2021). Not only the roots sense the hardness, but they also monitor the magnitude of hardness, as illustrated by a parallel increase in ROS and NO staining of root tips. In tomato, the increased NO levels in the root tip lead to inhibition of primary root elongation (Negi et al., 2010). Both ROS and NO have also been implicated in the initiation of lateral roots and root hairs (Tsukagoshi, 2016; Mur et al., 2013; Lombardo and Lamattina, 2012). Since hardness stimulated lateral root initiation and root hair formation in both VS and HS, it is plausible that these responses are interlinked with ROS and NO accumulation. Altogether, it can be surmised that hardness-imposed morphological responses appear to be causally related to stress signalling,

It is a general view that gravity-induced amyloplasts sedimentation in root columella cells leads to redistribution of auxin in root tip cells, which can be visualized by auxin sensitive reporters such as *IAA2::GUS* (Su, Gibbs, Jancewicz and Masson, 2017). In the HS root tip, which is presumably under incessant gravitational stress, the *IAA2::GUS* staining was nearly the same irrespective of hardness level. Likewise, in VS root tips too, the *IAA2::GUS* staining was similar. Evidently, the reduction of amyloplast staining or higher hardness levels did not affect auxin accumulation, at least in the root tip. Nonetheless, hardness did trigger morphological responses, wherein multiple hormones additively, synergistically, or antagonistically may play key roles.

Ethylene is one such hormone that affects several growth responses, including hardness-mediated inhibition of root elongation. Arabidopsis *ein2-1* mutant, which is compromised in ethylene signalling, does not display hardness-imposed inhibition of primary root elongation. Conversely, hardness-imposed inhibition of root elongation can be alleviated by ethylene action inhibitors (Okamoto et al., 2008). It was surmised that an alteration in ethylene signalling led to reduced root growth caused by horizontal hardness. While Okamoto et al. (2008) could not measure ethylene emission from wild-type seedlings, the ethylene emission from *eto1-1* mutant seedlings was nearly similar irrespective of horizontal or vertical orientation. Contrary to this, in tomato, the hardness stimulated ethylene emission from HS, whereas ethylene emission was not affected in VS. In our study, the usages of whole seedlings precluded determination of whether the ethylene specifically emanated from roots or shoots. In tomato, higher soil compaction stimulated ethylene emission from leaves; however, any root-specific ethylene emission was not determined (Roberts et al., 2002). Though the primary root and hypocotyl elongation were inhibited in both VS and HS, the above phenotypes were not causally linked, at least to ethylene emission.

In several growth responses, ethylene acts in a cohort with auxin. In tomato, the root penetration in soil requires coaction of ethylene and auxin (Santisree et al., 2011). Conversely, in seed germination, gibberellic acid and ABA act antagonistically (Liu and Hou, 2018). However, many developmental responses are not limited to specific hormone sets; most involve interaction among many hormones, including other regulatory molecules. The broad-spectrum analysis of plant hormones in roots and shoots of tomato seedlings with varying hardness levels is consistent with this view. First, hormonal levels are differentially affected in roots and shoots of seedlings. Second, the influence of hardness on individual hormones in roots and shoots differs in VS and HS. Lastly, hardness is likely perceived as stress, as evinced by major changes in hormones related to stress signalling.

The general view that root growth is influenced by auxin/ethylene balance seems not to be the only reason. Though HS seedlings emit more ethylene, the total auxin level in roots does not considerably decline. Conversely, IAA levels in VS roots decline continually, whereas ethylene emission is nearly similar. The near-uniform staining pattern of *IAA2::GUS* in root tips, though ethylene emission considerably increased in HS, also consonant to the above view. Unlike tomato seed germination, where a balance between ethylene and auxin facilitates the penetration of root tips in soil (Santisree et al., 2011), such a relationship is not apparent in hardness sensing.

Several studies have pointed out that both ABA and ethylene may act as signalling molecules when roots encounter compact soil. Soil compaction increases ABA levels in xylem sap, stimulates ethylene emission from tomato leaves (Roberts et al., 2002; Tracy et al., 2015). Since tomato ABA-deficient *notabilis* mutant emits higher ethylene, the ABA may be negatively regulating ethylene synthesis (Roberts et al., 2002). In our study, while VS does not show enhanced ethylene emission, it manifests several-fold higher ABA levels in roots and shoots. Conversely, in HS, the ABA increase in HS root and shoot is subdued than VS, but ethylene emission is several-fold stimulated in HS. Taken together, it is unlikely that ABA/ethylene coaction may be the reason for the VS and HS to show distinct phenotypes of roots/shoots of VS and HS.

Ostensibly hardness-induced morphological changes involve more complex interactions between different hormones. In Arabidopsis, hypoxia inhibits root growth by activating jasmonate signalling (Shukla et al., 2020). Since the seedlings had normal photoautotrophic phenotype, hypoxia-induced JA is least likely. JA application inhibits Arabidopsis’s primary root growth and stimulates the root hair and lateral root formation (Cai et al., 2014; Sun et al., 2009; Zhu et al., 2011). Given that JA levels increase only in VS roots, it negates JA’s role in the inhibition of root elongation. Treatment with exogenous SA inhibits Arabidopsis primary root growth and lateral root development (Pasternak et al., 2019). However, in our study, the suppression of primary root growth is associated with lateral root development. Since SA level declines on imposing hardness, it may not have a relationship with suppression of root growth and lateral root development in tomato.

Unlike the morphological responses that were similarly affected, the underlying metabolome was distinctly different in HS and VS. The metabolite alterations with increasing hardness and reduced growth indicate adaptation to sustain growth under stress conditions. Consistent with this, PCA of VS and HS metabolome revealed greater divergence in root and shoot with increasing hardness. The shift in the metabolic network at different hardness levels, too, reflects the above divergence. The increasing negative interaction in root networks points to the rewiring of metabolic networks to cope with the hardness. The preponderance of the negative interactions indicates that the hardness mediated metabolite shifts are designed to lower the metabolic fluxes across different pathways to cope up with the stress in roots.

The distinct difference, where amino acids are upregulated in horizontal roots, and sugars are upregulated in vertical roots, indicates that though hardness may be perceived as a common signal, the signal chains are divergent after perception. Distinct PCA profiles also corroborate the divergence in signalling. It can be construed that higher amino acids may arise by reduced protein synthesis, enhanced protein turnover, or increased biosynthesis (Huang and Jander, 2017). Notably, some amino acids are also are precursors for secondary metabolites and hormones. In HS, higher emission of ethylene is consistent with increased methionine levels. The contrasting upregulation of sugars and amino acids seems to be related to sustaining a balanced C/N ratio in impeded seedlings.

The commonly affected four metabolites viz. trehalose, dopamine, caffeoylquinic acid, and suberic acid seems to be the markers for the hardness stress. The altered levels of caffeoylquinic acid, an intermediate in the lignin biosynthesis pathway (e Silva, Mazzafera and Cesarino, 2019), and suberin, an effective apoplastic barrier (Vishwanath, Delude, Domergue and Rowland, 2015), seems to be related to cell wall strengthening in response to hardness. Under abiotic stress, lignin accumulation is induced, and its polymers are deposited in cell walls (Srivastava, Vishwakarma, Arafat, Gupta and Khan, 2015; Xu et al., 2020). The increased level of suberin with hardness is consistent with other stress-induced suberization such as salt and drought stress (Byrt, Munns, Burton, Gilliham and Wege, 2018; Krishnamurthy et al., 2009, Krishnamurthy, Ranathunge, Nayak, Schreiber and Mathew 2011). Ostensibly, the suberization and lignification of both roots and shoots are modulated by the magnitude of hardness.

In potato leaves, abiotic stress such as drought and UV radiation enhances dopamine levels (Świędrych, Stachowiak and Szopa, 2004). While little is known about the endogenous dopamine function, its exogenous application alleviates abiotic stress imposed by drought and salinity in apple seedlings (Li et al., 2015; Liang et al., 2018a). Considering that exogenous dopamine inhibits root elongation in soybean seedlings (Guidotti, Gomes, Siqueira-Soares, Soares and Ferrarese-Filho, 2013), the shortening of roots due to increased hardness may be causally related to the dopamine levels. Alternatively, high dopamine may ameliorate stress imposed by ROS by acting as an antioxidant, similar to that observed with dopamine-treated soybean and apple roots (Gomes et al., 2014; Jiao et al., 2019).

In contrast to dopamine, trehalose is a well-recognized stress ameliorator; its endogenous level is upregulated by various abiotic stresses such as temperature, salinity, and drought (Kosar, Akram, Sadiq, Al-Qurainy and Ashraf, 2019). Increased level of trehalose along with ABA may have bearing on inhibition of root growth, as trehalose and ABA synergistically inhibits Arabidopsis root growth (Wang et al., 2020). The roots of transgenic tomato seedlings with enhanced trehalose accumulation overcame exogenously-imposed oxidative stress more effectively than wild-type (Cortina and Culianez-Macia, 2005). Additionally, the increased trehalose, an osmotically compatible solute, can allow cells to build more pressure to sustain the root growth in compact media (Atwell, 1989).

The molecular mechanisms involved in hardness sensing are unknown; however, hardness affects several plasma membrane-localized proteins (Sampathkumar, 2020). The sensing of hardness by root may involve plasma membrane-localized mechanosensitive calcium channel MCA1 (Nakagawa et al., 2007), as mutants defective in MCA fails to penetrate hard agar surface (Denness et al., 2011). In such a case, hardness should elevate *MCA1* expression, especially in HS roots. Contrarily, at higher hardness, *MCA1* expression is lowered in HS and VS roots, thus discounting its role.

In general, post-germination, the growth is prioritized, and innate immunity is reduced (Lozano-Durán and Zipfel, 2015). The reduced expression of *RGS1,* a G-protein signaling effector, in VS and HS shoot may be related to a reduction in innate immunity response (Liang et al., 2018b) due to hardness-imposed altered root growth. The reduction in VS/HS growth also implies a lower frequency of cell divisions, which is related to cell cycle regulation. Consistent with this, the expression of *E2Fa,* a transcription factor regulating the onset of the S-phase of the cell cycle, is lowered in both VS/HS root and shoots. Interorgan communication also seems to play an important role in regulating growth responses. *HY5* is one such gene that affects multiple signalling pathways, including shoot to root communication. The hardness-triggered decline in *HY5* transcripts, in turn, may be causally linked with the initiation of more lateral roots (Sibout et al., 2006). The lower expression of *MYB36* at the higher hardness in VS and HS roots is consistent with the cessation of lateral root initiation at these hardness levels (Fernández-Marcos et al., 2017).

The relationship between the metabolome and gene expression is complex, as downstream to gene expression, the post-transcriptional, post-translation, and feed-forward/feed-back regulations of enzyme activities add to the above complexity. The reduced expression of *SUC trans (SUT1),* the major sucrose transporter in the phloem (Hackel et al. 2006), correlates with sugar accumulation in VS and HS shoots, but not with sugar accumulation in VS root, which likely depends on sugar derived from the shoot. Little correlation with sugar accumulation was observed in the expression of other sugar-related genes *HXK1*, *Lin6*, *SUS3*, and *invertases.* Considering that VS shoot has low *LIN6* expression but has a higher accumulation of fructose-1-phosphate, it is consistent with the above view.

The accumulation of trehalose in VS and HS seems to indicate its role other than that of stress ameliorator. Since overaccumulation of trehalose may drastically affect the plant’s C/N balance, reduced expression of *TRE2* in VS/HS roots at higher hardness may be related to maintaining the above balance. Additionally, trehalose-6-phosphate, the precursor of trehalose, inhibits the activity of the SnRK1 (Baena-González and Lunn, 2020; Paul, Watson and Griffiths, 2020; Zhang et al., 2009) and may balance growth versus stress response by modulating RAPTOR phosphorylation (Rosenberger and Chen, 2018; Wang et al., 2018). The higher accumulation of trehalose indicates a higher level of its precursor trehalose-6-phosphate, which may lower the *SnRK1* expression (Fichtner and Lunn, 2021). Consistent with this, the *SnRK1a* expression is subdued in both VS/HS root and shoot at all hardness levels.

The metabolic homeostasis of cells is also maintained by TOR/RAPTOR signalling modules. Contrary to *SnRK1,* the TOR/RAPTOR signalling may be affected only beyond ? 1% hardness level, where the expression of *TEL, RAPTOR, Lst8,* and *S6K,* the key genes of the above signalling pathway, are lowered. Though trehalose is a signature metabolite, it is equally plausible that the shift in the levels of phytohormones, IAA (Retzer and Weckwerth, 2021), and ABA (Belda-Palazón et al., 2020; Rodrigues et al., 2013) can also modulate the TOR and SnRK signalling pathway. Ostensibly, the interrelationship between phytohormones levels, metabolic homeostasis, SnRK1, and TOR/RAPTOR pathway is complex. Between, SnRK1 and TOR/RAPTOR signalling pathways, the SnRK1 seems to be the major contender. Notwithstanding the above, the observed correlations are only indicative but can be explored in future studies.

As the plant life cycle begins under the soil cover, the radicle right after emergence from seed encounters hardness, sensing which modulates growth of both shoot and root. Our study highlights that hardness imposition affects a wide range of responses encompassing redox signalling, phytohormone levels, and metabolome shifts. Though these responses are triggered by hardness, the interrelationship between these responses remains to be deciphered. An important question is whether the root tip senses the hardness or the entire organ can sense it. Though many plasma-membrane localized proteins are affected by mechanical stress, none of these have emerged as a key sensor. The uniform *IAA2::GUS* staining in VS and HS root tips at all hardness levels indicates that unlike directional stimuli such as gravity, the hardness is likely sensed beyond the root tip, akin to hydrotropism (Dietrich et al., 2017). In the future, the knowledge about hardness sensing by a receptor or a cohort of regulators would help to uncover signalling emanating from hardness sensing leading to a shift in metabolic homeostasis.

## Data Availability

All data associated with this manuscript is provided in supplemental data.

## Author Contributions

R.S and Y.S. designed this project and R.S, Y.S and A.K. wrote the manuscript. A.K. performed most of the experiments. S.N. did the microscope imaging. S.D. made the correlation networks.

## Supplemental Information

Supplemental tables and figures are available online.

## Supporting data

**Figure S1:** Horizontal/vertical oriented seedling growth setup and phenotype of seedlings.

**Figure S2:** Reduction in epidermal cell length of primary roots of horizontally/vertically-impeded seedlings.

**Figure S3:** Effect of hardness on primary root diameter.

**Figure S4:** Effect of hardness on the expression of auxin reporter *IAA2::GUS* in different organs of seedling

**Figure S5:** Impact of increasing hardness on metabolite/hormone correlation networks.

**Figure S6:** Reduction in shared interaction among metabolites with increasing hardness.

**Table S1**: List of genes and the primers used for qRT-PCR analysis.

**Table S2:** The hormones detected in roots and shoots with log 2 and *p* value.

**Table S3:** The metabolites detected in roots with log 2 and *p* value.

**Table S4:** The metabolites detected in shoots with log 2 and *p* value.

**Table S5:** Unique and shared interactions in metabolic networks of root and shoot.

**Table S6:** The transcripts levels of C/N metabolism genes in roots and shoots.

## References

Arnaud, C., Bonnot, C., Desnos, T., & Nussaume, L. (2010). The root cap at the forefront. Comptes Rendus Biologies, 333, 335–343.

Atwell, B. J. (1989). Physiological responses of lupin roots to soil compaction. In Structural and Functional Aspects of Transport in Roots. (eds B.C. Loughman, O. Gasparíková, & J. Kolek), (pp. 251–255). Springer, Dordrecht. https://doi.org/10.1007/BF02139953.

Belda-Palazón, B., Adamo, M., Valerio, C., Ferreira, L. J., Confraria, A., Reis-Barata, D., Rodrigues, A., Meyer, C., Rodriguez, P. L. & Baena-González, E., (2020). A dual function of SnRK2 kinases in the regulation of SnRK1 and plant growth. Nature Plants, 6, 1345–1353.

Baena-González, E. and Lunn, J.E., (2020). SnRK1 and trehalose 6-phosphate–two ancient pathways converge to regulate plant metabolism and growth. Current Opinion in Plant Biology, 55, 52–59.

Benjamins, R., Quint, A. B., Weijers, D., Hooykaas, P., & Offringa, R. (2001). The PINOID protein kinase regulates organ development in Arabidopsis by enhancing polar auxin transport. Development, 128, 4057–4067.

Byrt, C. S., Munns, R., Burton, R. A., Gilliham, M., & Wege, S. (2018). Root cell wall solutions for crop plants in saline soils. Plant Science, 269, 47–55.

Cai, X. T., Xu, P., Zhao, P. X., Liu, R., Yu, L. H., & Xiang, C. B. (2014). Arabidopsis ERF109 mediates cross-talk between jasmonic acid and auxin biosynthesis during lateral root formation. Nature Communications, 5, 1–13.

Cortina, C., & Culiáñez-Macià, F. A. (2005). Tomato abiotic stress enhanced tolerance by trehalose biosynthesis. Plant Science, 169, 75–82.

Denness, L., McKenna, J. F., Segonzac, C., Wormit, A., Madhou, P., Bennett, M., Mansfield, J., Zipfel, C. & Hamann, T. (2011). Cell wall damage-induced lignin biosynthesis is regulated by a reactive oxygen species-and jasmonic acid-dependent process in Arabidopsis. Plant Physiology, 156, 1364–1374.

Dietrich, D., Pang, L., Kobayashi, A., Fozard, J.A., Boudolf, V., Bhosale, R., et al. (2017). Root hydrotropism is controlled via a cortex-specific growth mechanism. Nature plants, 3, 1–8.

e Silva, N. V., Mazzafera, P., & Cesarino, I. (2019). Should I stay or should I go: are chlorogenic acids mobilized towards lignin biosynthesis? Phytochemistry, 166, 112063.

Farnese, F. S., Menezes-Silva, P. E., Gusman, G. S., & Oliveira, J. A. (2016). When bad guys become good ones: the key role of reactive oxygen species and nitric oxide in the plant responses to abiotic stress. Frontiers in Plant Science, 7, 471.

Fernández-Marcos, M., Desvoyes, B., Manzano, C., Liberman, L. M., Benfey, P. N., del Pozo, J. C., & Gutierrez, C. (2017). Control of Arabidopsis lateral root primordium boundaries by MYB 36. New Phytologist, 213, 105–112.

Fichtner, F. & Lunn, J.E., (2021). The role of trehalose 6-phosphate (Tre6P) in plant metabolism and development. Annual Review of Plant Biology, 72. https://doi.org/10.1146/annurev-arplant-050718-095929

Fruleux, A., Verger, S., & Boudaoud, A. (2019). Feeling stressed or strained? A biophysical model for cell wall mechanosensing in plants. Frontiers in Plant Science, 10, 757.

Gomes, B. R., Siqueira-Soares, R. D. C., Santos, W. D. D., Marchiosi, R., Soares, A. R., & Ferrarese-Filho, O. (2014). The effects of dopamine on antioxidant enzymes activities and reactive oxygen species levels in soybean roots. Plant Signaling & Behavior, 9, e977704. https://doi.org/10.4161/15592324.2014.977704

Guidotti, B. B., Gomes, B. R., Siqueira-Soares, R. D. C., Soares, A. R., & Ferrarese-Filho, O. (2013). The effects of dopamine on root growth and enzyme activity in soybean seedlings. Plant Signaling & Behavior, 8, e25477. https://dx.doi.org/10.4161/psb.25477

Hackel, A., Schauer, N., Carrari, F., Fernie, A. R., Grimm, B., & Kühn, C. (2006). Sucrose transporter LeSUT1 and LeSUT2 inhibition affects tomato fruit development in different ways. The Plant Journal, 45, 180–192.

Hanbury, C. D., & Atwell, B. J. (2005). Growth dynamics of mechanically impeded lupin roots: does altered morphology induce hypoxia? Annals of Botany, 96, 913–924.

Hashiguchi, Y., Tasaka, M., & Morita, M. T. (2013). Mechanism of higher plant gravity sensing. American Journal of Botany, 100, 91–100.

Hildreth, S. B., Foley, E. E., Muday, G. K., Helm, R. F., & Winkel, B. S. (2020). The dynamic response of the Arabidopsis root metabolome to auxin and ethylene is not predicted by changes in the transcriptome. Scientific Reports, 10, 1–15.

Huang, T., & Jander, G. (2017). Abscisic acid-regulated protein degradation causes osmotic stress-induced accumulation of branched-chain amino acids in Arabidopsis thaliana. Planta, 246, 737–747.

Iijima, M., Griffiths, B., & Bengough, A. G. (2000). Sloughing of cap cells and carbon exudation from maize seedling roots in compacted sand. New Phytologist, 145, 477–482.

Jiao, X., Li, Y., Zhang, X., Liu, C., Liang, W., Li, C., Ma, F. & Li, C. (2019). Exogenous Dopamine Application Promotes Alkali Tolerance of Apple Seedlings. Plants, 8, 580.

Jacobsen, A. G. R., Jervis, G., Xu, J., Topping, J. F. & Lindsey, K. (2021) Root growth responses to mechanical impedance are regulated by a network of ROS, ethylene and auxin signalling in Arabidopsis. New Phytologist, https://doi.org/10.1111/nph.17180

Kosar, F., Akram, N. A., Sadiq, M., Al-Qurainy, F., & Ashraf, M. (2019). Trehalose: a key organic osmolyte effectively involved in plant abiotic stress tolerance. Journal of Plant Growth Regulation, 38, 606–618.

Krishnamurthy, P., Ranathunge, K., Franke, R., Prakash, H. S., Schreiber, L., & Mathew, M. K. (2009). The role of root apoplastic transport barriers in salt tolerance of rice (Oryza sativa L.). Planta, 230, 119–134.

Krishnamurthy, P., Ranathunge, K., Nayak, S., Schreiber, L., & Mathew, M. K. (2011). Root apoplastic barriers block Na+ transport to shoots in rice (Oryza sativa L.). Journal of Experimental Botany, 62, 4215–4228.

Kumari, A., Ray, K., Sadhna, S., Pandey, A. K., Sreelakshmi, Y., & Sharma, R. (2017). Metabolomic homeostasis shifts after callus formation and shoot regeneration in tomato. PloS one, 12, e0176978.

Lee, H. J., Kim, H. S., Park, J. M., Cho, H. S., & Jeon, J. H. (2020). PIN-mediated polar auxin transport facilitates root-obstacle avoidance. New Phytologist, 225, 1285–1296.

Li, C., Sun, X., Chang, C., Jia, D., Wei, Z., Li, C., & Ma, F. (2015). Dopamine alleviates salt-induced stress in Malus hupehensis. Physiologia Plantarum, 153, 584–602.

Liang, B., Gao, T., Zhao, Q., Ma, C., Chen, Q., Wei, Z., Li, C., Li, C. & Ma, F. (2018a). Effects of exogenous dopamine on the uptake, transport, and resorption of apple ionome under moderate drought. Frontiers in Plant Science, 9, 755.

Liang, X., Ma, M., Zhou, Z., Wang, J., Yang, X., Rao, S., Bi, G., Li, L., Zhang, X., Chai, J. & Zhou, J. M. (2018b). Ligand-triggered de-repression of Arabidopsis heterotrimeric G proteins coupled to immune receptor kinases. Cell Research, 28, 529–543.

Liu, X., & Hou, X. (2018). Antagonistic regulation of ABA and GA in metabolism and signaling pathways. Frontiers in Plant Science, 9, 251.

Lombardo, M. C., & Lamattina, L. (2012). Nitric oxide is essential for vesicle formation and trafficking in Arabidopsis root hair growth. Journal of Experimental Botany, 63, 4875–4885.

Lozano-Durán, R., & Zipfel, C. (2015). Trade-off between growth and immunity: role of brassinosteroids. Trends in Plant Science, 20, 12–19.

Moss, G. I., Hall, K. C., & Jackson, M. B. (1988). Ethylene and the responses of roots of maize (Zea mays L.) to physical impedance. New Phytologist, 109, 303–311.

Mousavi, S. A. R., Dubin, A. E., Zeng, W. Z., Coombs, A. M., Do, K., Ghadiri, D. A., Ge, C., Ge, C., Zhao, Y. & Patapoutian, A., (2020). PIEZO ion channel is required for root mechanotransduction in Arabidopsis thaliana. BioRxiv, https://doi.org/10.1101/2020.08.27.270355

Mur, L. A., Mandon, J., Persijn, S., Cristescu, S. M., Moshkov, I. E., Novikova, G. V., Hall, M.A., Harren, F. J., Hebelstrup, K. H. & Gupta, K. J. (2013). Nitric oxide in plants: an assessment of the current state of knowledge. AoB plants, 5, pls052

Nakagawa, Y., Katagiri, T., Shinozaki, K., Qi, Z., Tatsumi, H., Furuichi, T., Kishigami, A., Sokabe, M., Kojima, I., Sato, S. and Kato, T., & Iida, H. (2007). Arabidopsis plasma membrane protein crucial for Ca2+ influx and touch sensing in roots. Proceedings of the National Academy of Sciences, 104, 3639–3644.

Negi, S., Santisree, P., Kharshiing, E. V., & Sharma, R. (2010). Inhibition of the ubiquitin— proteasome pathway alters cellular levels of nitric oxide in tomato seedlings. Molecular Plant, 3, 854–869.

Okamoto, K., & Yano, K. (2017). Al resistance and mechanical impedance to roots in Zea mays: Reduced Al toxicity via enhanced mucilage production. Rhizosphere, 3, 60–66.

Okamoto, T., Tsurumi, S., Shibasaki, K., Obana, Y., Takaji, H., Oono, Y., & Rahman, A. (2008). Genetic dissection of hormonal responses in the roots of Arabidopsis grown under continuous mechanical impedance. Plant Physiology, 146, 1651–1662.

Oyama, T., Shimura, Y., & Okada, K. (1997). The Arabidopsis HY5 gene encodes a bZIP protein that regulates stimulus-induced development of root and hypocotyl. Genes & Development, 11, 2983–2995.

Pan, X., Welti, R., & Wang, X. (2010). Quantitative analysis of major plant hormones in crude plant extracts by high-performance liquid chromatography–mass spectrometry. Nature Protocols, 5, 986–992.

Pandey, B. K., Huang, G., Bhosale, R., Hartman, S., Sturrock, C. J., Jose, L., et al. (2021). Plant roots sense soil compaction through restricted ethylene diffusion. Science, 371, 276–280.

Pasternak, T., Groot, E. P., Kazantsev, F. V., Teale, W., Omelyanchuk, N., Kovrizhnykh, V., Palme, K. & Mironova, V. V. (2019). Salicylic acid affects root meristem patterning via auxin distribution in a concentration-dependent manner. Plant Physiology, 180, 1725–1739.

Paul, M. J., Watson, A., & Griffiths, C. A. (2020). Trehalose 6-phosphate signalling and impact on crop yield. Biochemical Society Transactions, 48, 2127–2137.

Pelagio-Flores, R., Ruiz-Herrera, L. F., & López-Bucio, J. (2016). Serotonin modulates Arabidopsis root growth via changes in reactive oxygen species and jasmonic acid–ethylene signaling. Physiologia Plantarum, 158, 92–105.

Potocka, I., & Szymanowska-Pułka, J. (2018). Morphological responses of plant roots to mechanical stress. Annals of Botany, 122, 711–723.

Retzer, K. & Weckwerth, W. (2021). The TOR–Auxin Connection Upstream of Root Hair Growth. Plants, 10, 150. https://doi.org/10.3390/plants10010150

Roberts, J. A., Hussain, A., Taylor, I. B., & Black, C. R. (2002). Use of mutants to study long-distance signalling in response to compacted soil. Journal of Experimental Botany, 53, 45–50.

Roessner, U., Wagner, C., Kopka, J., Trethewey, R. N., & Willmitzer, L. (2000). Simultaneous analysis of metabolites in potato tuber by gas chromatography–mass spectrometry. The Plant Journal, 23, 131–142.

Rodrigues, A., Adamo, M., Crozet, P., Margalha, L., Confraria, A., Martinho, C., Elias, A., Rabissi, A., Lumbreras, V., González-Guzmán, M. & Antoni, R., (2013) ABI1 and PP2CA phosphatases are negative regulators of snf1-related protein kinase1 signaling in Arabidopsis. Plant Cell, 25, 3871–3884.

Rosenberger, C. L., & Chen, J. (2018). To grow or not to grow: TOR and SnRK2 coordinate growth and stress response in Arabidopsis. Molecular Cell, 69, 3–4.

Růžička, K., Ljung, K., Vanneste, S., Podhorská, R., Beeckman, T., Friml, J., & Benková, E. (2007). Ethylene regulates root growth through effects on auxin biosynthesis and transport-dependent auxin distribution. The Plant Cell, 19, 2197–2212.

Sampathkumar, A. (2020). Mechanical feedback-loop regulation of morphogenesis in plants. Development, 147, dev177964.

Santisree, P., Nongmaithem, S., Vasuki, H., Sreelakshmi, Y., Ivanchenko, M. G., & Sharma, R. (2011). Tomato root penetration in soil requires a coaction between ethylene and auxin signaling. Plant Physiology, 156, 1424–1438.

Sarquis, J. I., Jordan, W. R., & Morgan, P. W. (1991). Ethylene evolution from maize (Zea mays L.) seedling roots and shoots in response to mechanical impedance. Plant Physiology, 96, 1171–1177.

Schumacher, T. E., & Smucker, A. J. M. (1981). Mechanical Impedance Effects on Oxygen Uptake and Porosity of Dry bean Roots. Agronomy Journal, 73, 51–55.

Shih, H. W., Miller, N. D., Dai, C., Spalding, E. P., & Monshausen, G. B. (2014). The receptor-like kinase FERONIA is required for mechanical signal transduction in Arabidopsis seedlings. Current Biology, 24, 1887–1892.

Shukla, V., Lombardi, L., Pencik, A., Novak, O., Weits, D.A., Loreti, E., Perata, P., Giuntoli, B. & Licausi, F. (2020). Jasmonate signalling contributes to primary root inhibition upon oxygen deficiency in Arabidopsis thaliana. Plants, 9, 1046.

Sibout, R., Sukumar, P., Hettiarachchi, C., Holm, M., Muday, G. K., & Hardtke, C. S. (2006). Opposite root growth phenotypes of hy5 versus hy5 hyh mutants correlate with increased constitutive auxin signaling. PLoS Genet, 2, e202.

Souty, N., & Rode, C. (1987). Aspect mécanique de la croissance des racines. I.-Mesure de la force de pénétration. Agronomie, EDP Sciences, 7, 623–630.

Srivastava, S., Vishwakarma, R. K., Arafat, Y. A., Gupta, S. K., & Khan, B. M. (2015). Abiotic stress induces change in Cinnamoyl CoA Reductase (CCR) protein abundance and lignin deposition in developing seedlings of Leucaena leucocephala. Physiology and Molecular Biology of Plants, 21, 197–205.

Su, S. H., Gibbs, N. M., Jancewicz, A. L., & Masson, P. H. (2017). Molecular mechanisms of root gravitropism. Current Biology, 27, R964–R972.

Sun, J., Xu, Y., Ye, S., Jiang, H., Chen, Q., Liu, F., et al. (2009). Arabidopsis ASA1 is important for jasmonate-mediated regulation of auxin biosynthesis and transport during lateral root formation. The Plant Cell, 21, 1495–1511.

Swarup, R., Perry, P., Hagenbeek, D., Van Der Straeten, D., Beemster, G. T., Sandberg, G., Bhalerao, R., Ljung, K. & Bennett, M. J. (2007). Ethylene upregulates auxin biosynthesis in Arabidopsis seedlings to enhance inhibition of root cell elongation. The Plant Cell, 19, 2186–2196.

Świędrych, A., Stachowiak, J., & Szopa, J. (2004). The catecholamine potentiates starch mobilization in transgenic potato tubers. Plant Physiology and Biochemistry, 42, 103–109.

Takahashi, N., Yamazaki, Y., Kobayashi, A., Higashitani, A., & Takahashi, H. (2003). Hydrotropism interacts with gravitropism by degrading amyloplasts in seedling roots of Arabidopsis and radish. Plant Physiology, 132, 805–810.

Tracy, S. R., Black, C. R., Roberts, J. A., Dodd, I. C., & Mooney, S. J. (2015). Using X-ray computed tomography to explore the role of abscisic acid in moderating the impact of soil compaction on root system architecture. Environmental and Experimental Botany, 110, 11–18.

Tracy, S. R., Black, C. R., Roberts, J. A., Sturrock, C., Mairhofer, S., Craigon, J., & Mooney, S. J. (2012). Quantifying the impact of soil compaction on root system architecture in tomato (Solanum lycopersicum) by X-ray micro-computed tomography. Annals of Botany, 110, 511–519.

Tsukagoshi, H. (2016). Control of root growth and development by reactive oxygen species. Current Opinion in Plant Biology, 29, 57–63.

Vishwanath, S. J., Delude, C., Domergue, F., & Rowland, O. (2015). Suberin: biosynthesis, regulation, and polymer assembly of a protective extracellular barrier. Plant Cell Reports, 34, 573–586.

Wang, P., Zhao, Y., Li, Z., Hsu, C. C., Liu, X., Fu, L et al. (2018). Reciprocal regulation of the TOR kinase and ABA receptor balances plant growth and stress response. Molecular Cell, 69, 100–112.

Wang, W., Chen, Q., Xu, S., Liu, W. C., Zhu, X. & Song, C.P. (2020). Trehalose-6-phosphate phosphatase E modulates ABA-controlled root growth and stomatal movement in Arabidopsis. Journal of Integrative Plant Biology. 62, 1518–1534.

Wilson, A. J., & Robards, A. W. (1978). The ultrastructural development of mechanically impeded barley roots. Effects on the endodermis and pericycle. Protoplasma, 95, 255–265.

Wilson, A. J., Robards, A. W., & Goss, M. J. (1977). Effects of mechanical impedance on root growth in barley, Hordeum vulgare L. II. Effects on cell development in seminal roots. Journal of Experimental Botany, 28, 1216–1227.

Xu, W., Tang, W., Wang, C., Ge, L., Sun, J., Qi, X., et al. (2020). SiMYB56 confers drought stress tolerance in transgenic rice by regulating lignin biosynthesis and ABA signaling pathway. Frontiers in Plant Science, 11, 785.

Zhang Y, He P, Ma X, Yang Z, Pang C, Yu J, Wang G, Friml J, & Xiao G. (2019) Auxin-mediated statolith production for root gravitropism. New Phytologist, 224, 761–74.

Zhang, Y., Primavesi, L. F., Jhurreea, D., Andralojc, P. J., Mitchell, R. A., Powers, S. J., Schluepmann, H., Delatte, T., Wingler, A. & Paul, M. J. (2009). Inhibition of SNF1-related protein kinase1 activity and regulation of metabolic pathways by trehalose-6-phosphate. Plant Physiology, 149, 1860–1871.

Zhu, Z., An, F., Feng, Y., Li, P., Xue, L., Mu, A., et al. (2011). Derepression of ethylene-stabilized transcription factors (EIN3/EIL1) mediates jasmonate and ethylene signaling synergy in Arabidopsis. Proceedings of the National Academy of Sciences, 108, 12539–12544.

