## Supplementary material for "Tomato roots sense horizontal/vertical mechanical impedance and divergently modulate root/shoot metabolome": Figure S1

Supplementary Figure S1

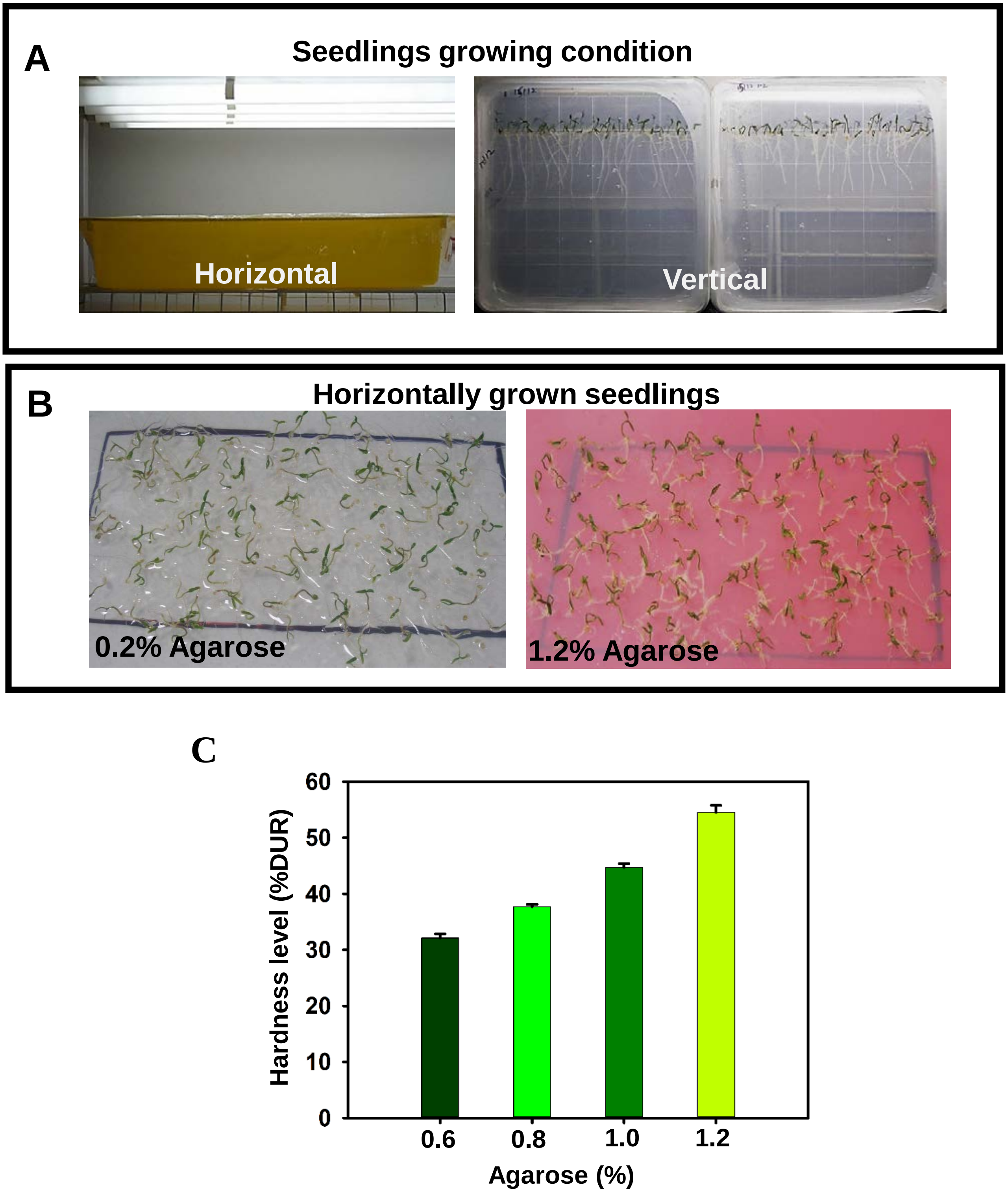

Supplementary Figure S1

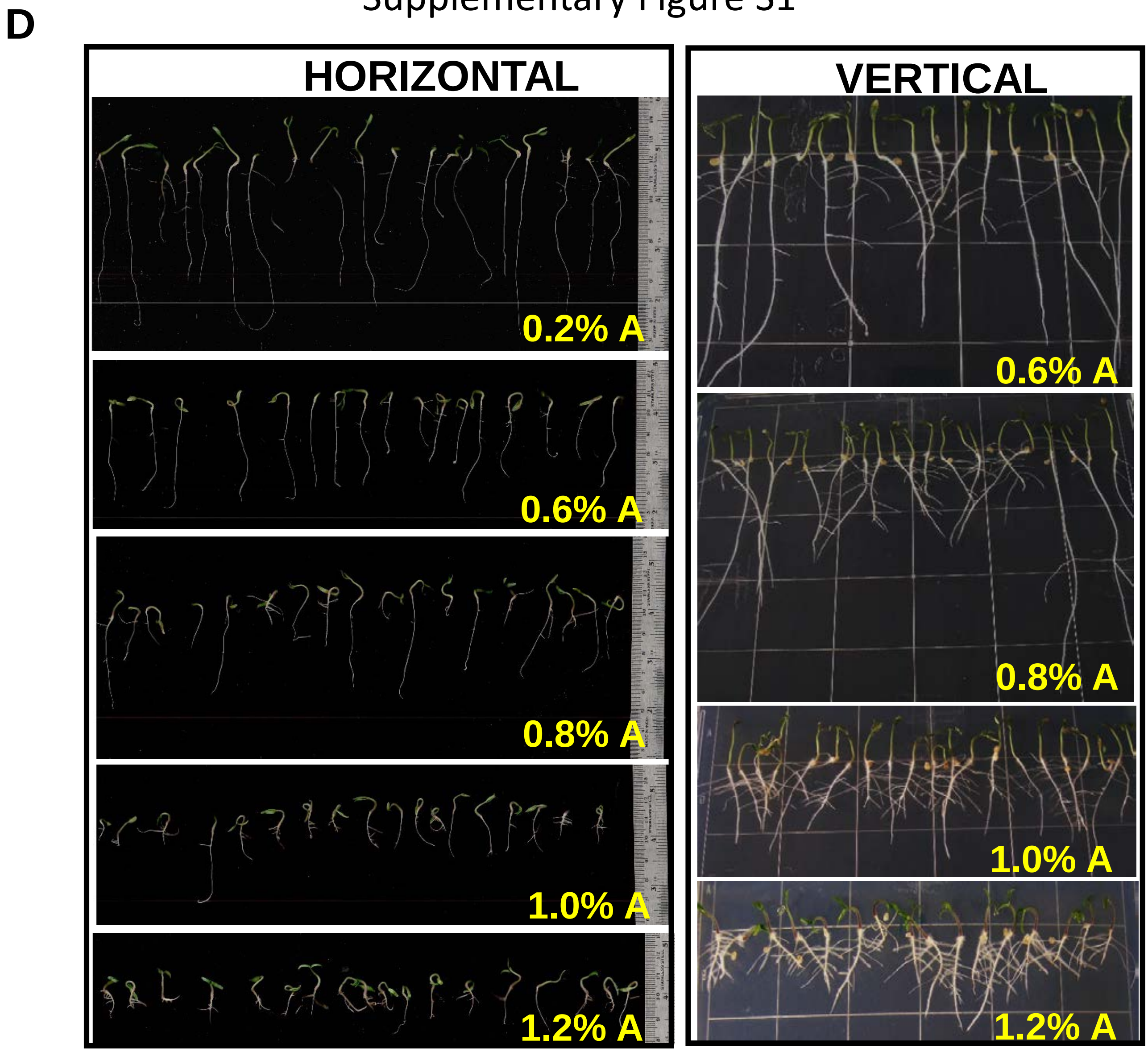


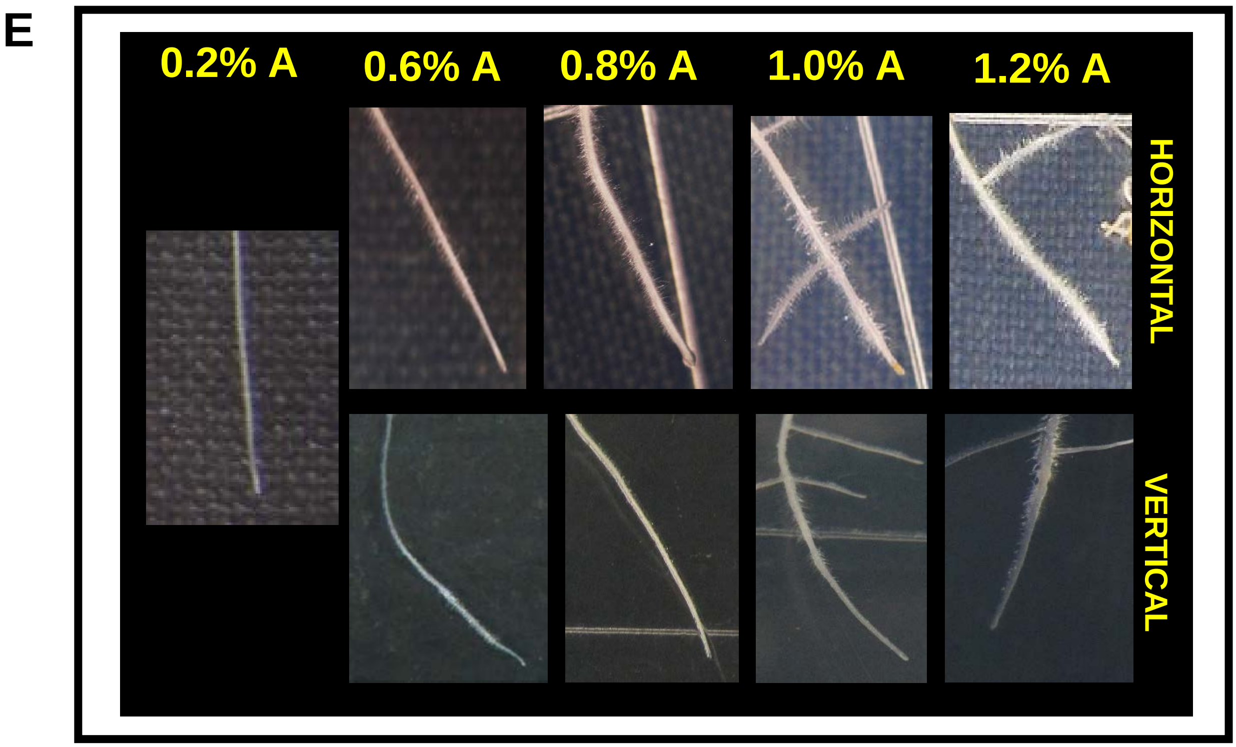

### Supplementary Figure S1

**Figure S1: Horizontal/vertical oriented seedling growth setup and phenotype of seedlings.**
**A,** The trays (Size 48.0 *l* × 36.0 *w* ×9 *h* cm) and plates (Size 24.0 *l* × 24.0 *w* ×1.2 *h* cm) used for growing tomato seedlings in horizontal and vertical orientation, respectively. **B,** Phenotype of seedlings grown horizontally at 0.2%A and 1.2%A. Note at 0.2%A, the roots can penetrate and grow within agarose layer. **C,** Progressive increase in agarose hardness with increasing concentration. The hardness was measured by using DUROFEL DFT 100 (Agrosta, France). **D,** Phenotype of seedlings grown vertically or horizontally on 0.2, 0.6, 0.8, 1.0 and 1.2% of agarose (w/v). **Note** progressive reduction in primary root length and increased proliferation of lateral roots with increasing hardness. **E,** Close up photographs of primary roots showing increase in root hair density and proximity to the root tip with increasing hardness. In all images **A** indicates Agarose unless otherwise mentioned.

Supplementary Figure S2

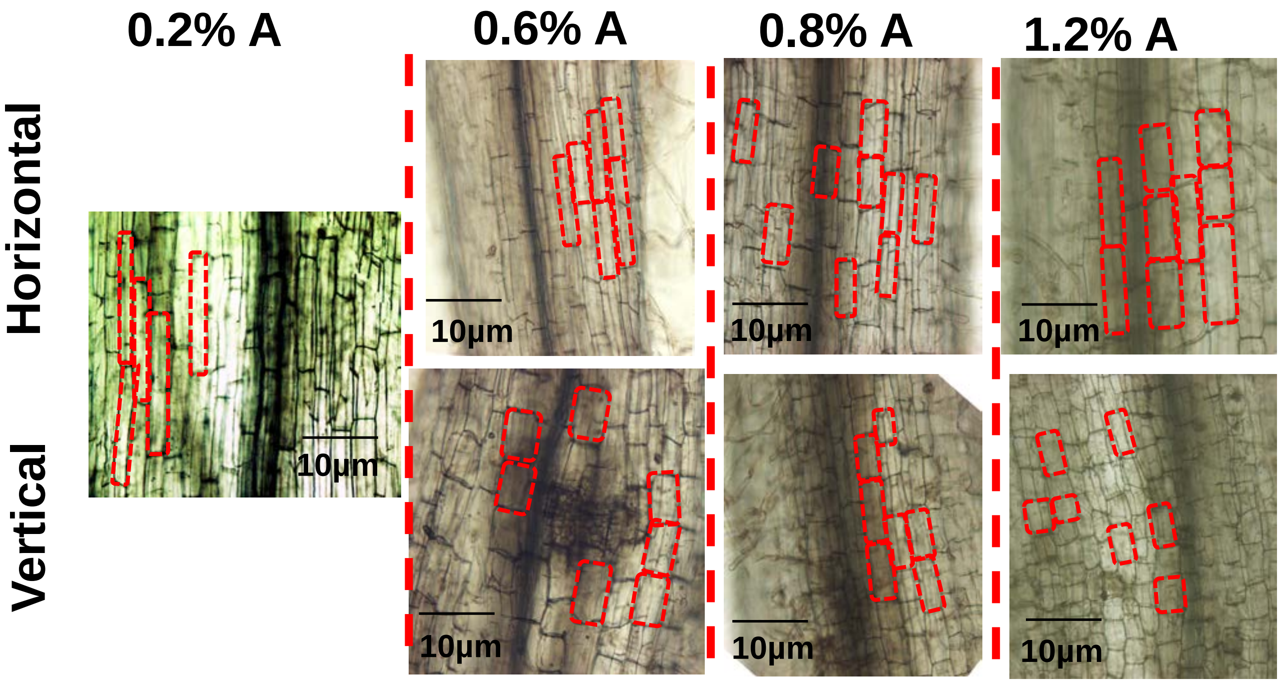


**Figure S2 : Reduction in epidermal cell length of primary roots of horizontally/vertically-impeded seedlings**.

The bright-field images of root tips were captured using an Olympus BX-45 fluorescent microscope. The region selected for comparing the cell length was approximately 10 mm distal from the root tip. To highlight the difference in cell lengths, the boundaries of 2-3 cell in each tip were marked with dotted red lines.

Supplementary Figure S3

A

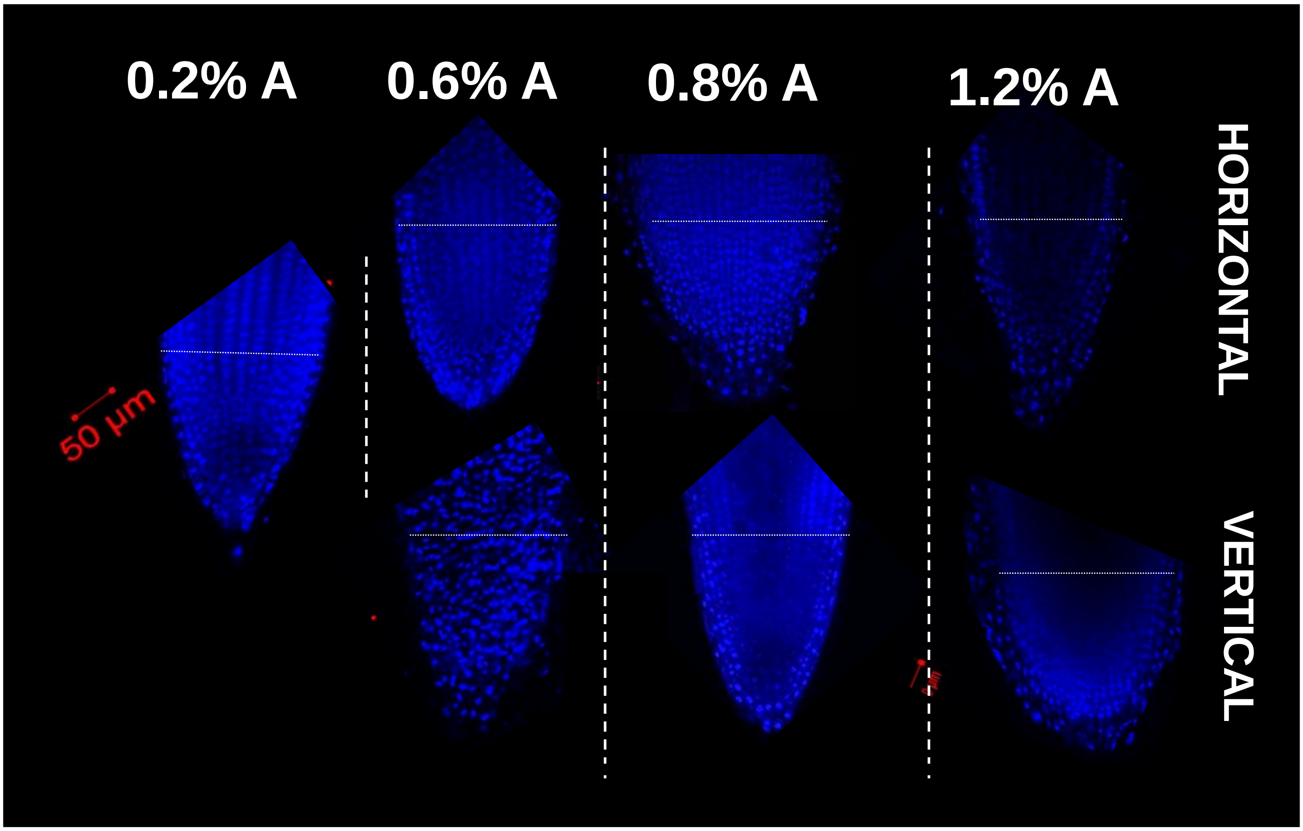


B

| Agarose(%) | Number of cell layers | |
|---|---|---|
| | Horizontally-impeded root | Vertically-impeded root |
| 0.2% | ~19 | -- |
| 0.6% | ~21 | ~20 |
| 0.8% | ~24 | ~21 |
| 1.2% | ~25 | ~20 |

C

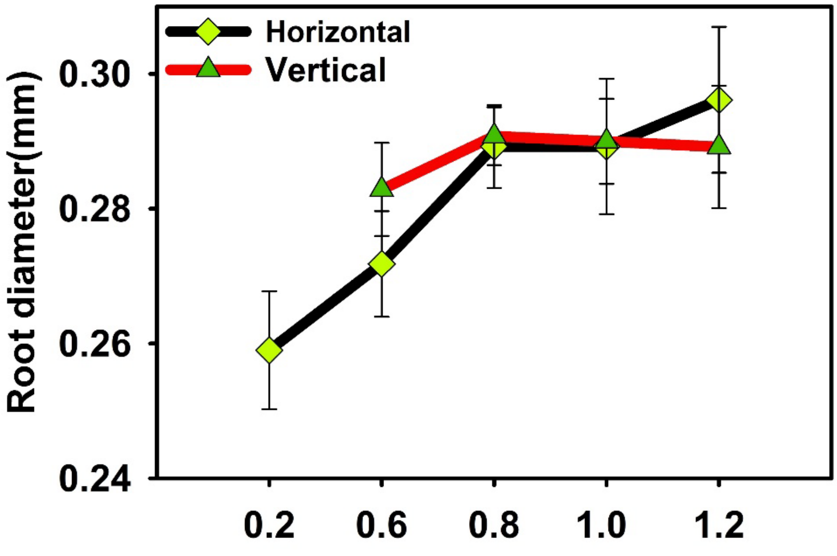

### Supplementary Figure S3

**Figure S3: Effect of hardness on primary root diameter.**
**A,** DAPI stained primary root-tips of horizontally and vertically impeded 5-day old seedlings. Images were captured using Zeiss confocal microscope, using 358 nm excitation and 461 nm emission wavelength. The horizontal white line overlaid on the image indicates the region selected for counting the cell layers. **B,** The number of cell layers counted across the primary root at different hardness. An increase in the number of cell layers was observed in the horizontal roots at higher hardness. The number of cell layers in vertically impeded root were nearly similar to those observed in M82 primary root sections [Ron et al. (2013) Identification of novel loci regulating interspecific variation in root morphology and cellular development in tomato. *Plant Physiology.* **162**:755-768]. **C,** Increase in root diameter of the seedlings with increasing hardness.

Supplementary Figure S4

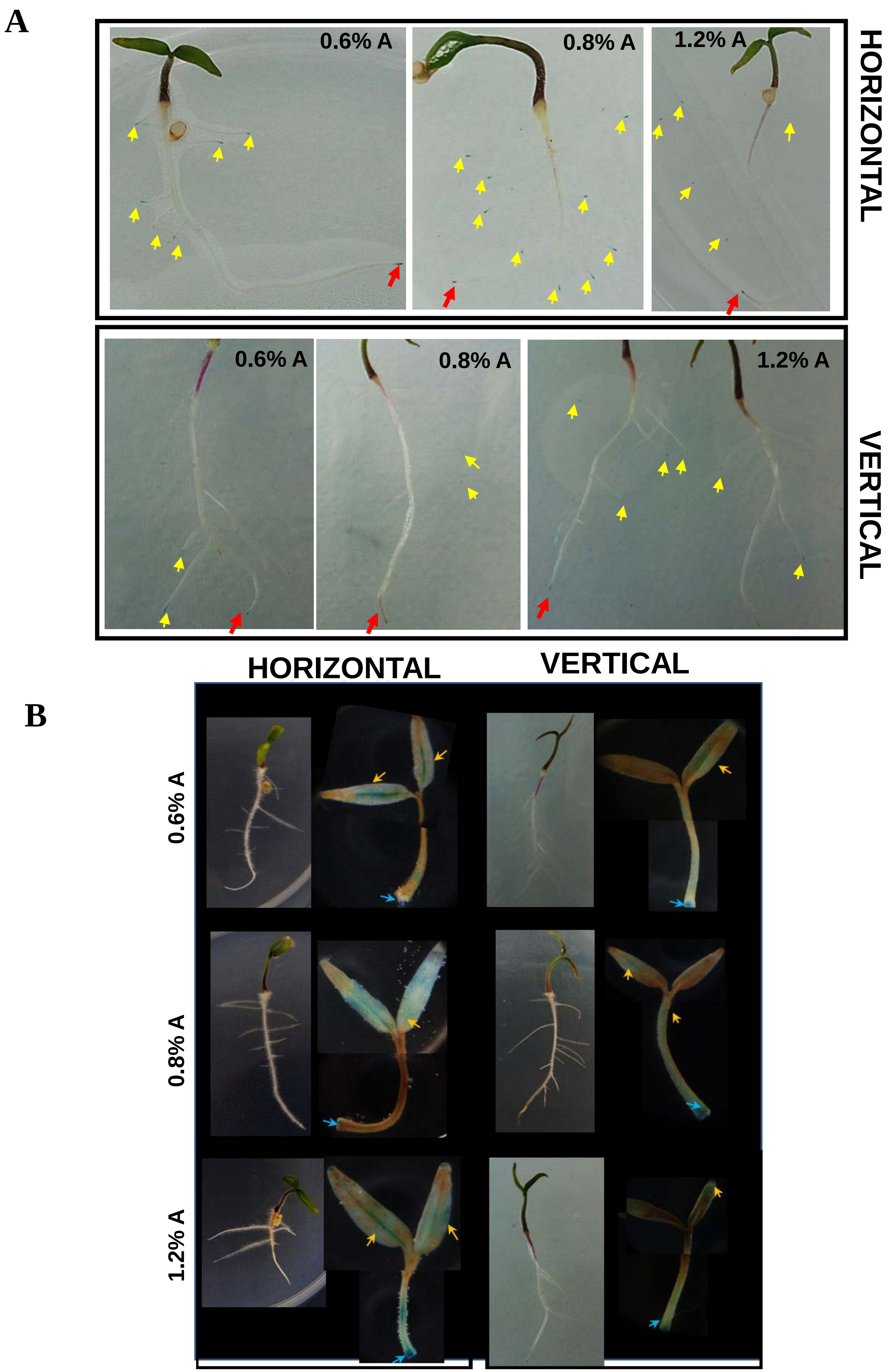

### Supplementary Figure S4

**Figure S4: Effect of hardness on the expression of auxin reporter *IAA2::GUS* in different organs of seedling.**

**A,** Seedling picture depicting GUS staining in primary/lateral root tips. **Note** the GUS staining is more intense in the horizontally impeded root tips than vertically impeded root. The yellow and the red colour arrow represents the main and lateral root respectively. **B**, The GUS staining of the whole shoot. The yellow arrow displays the GUS staining area in cotyledons whereas blue arrow indicates the GUS staining area in shoot and root junction.

Supplementary Figure S5

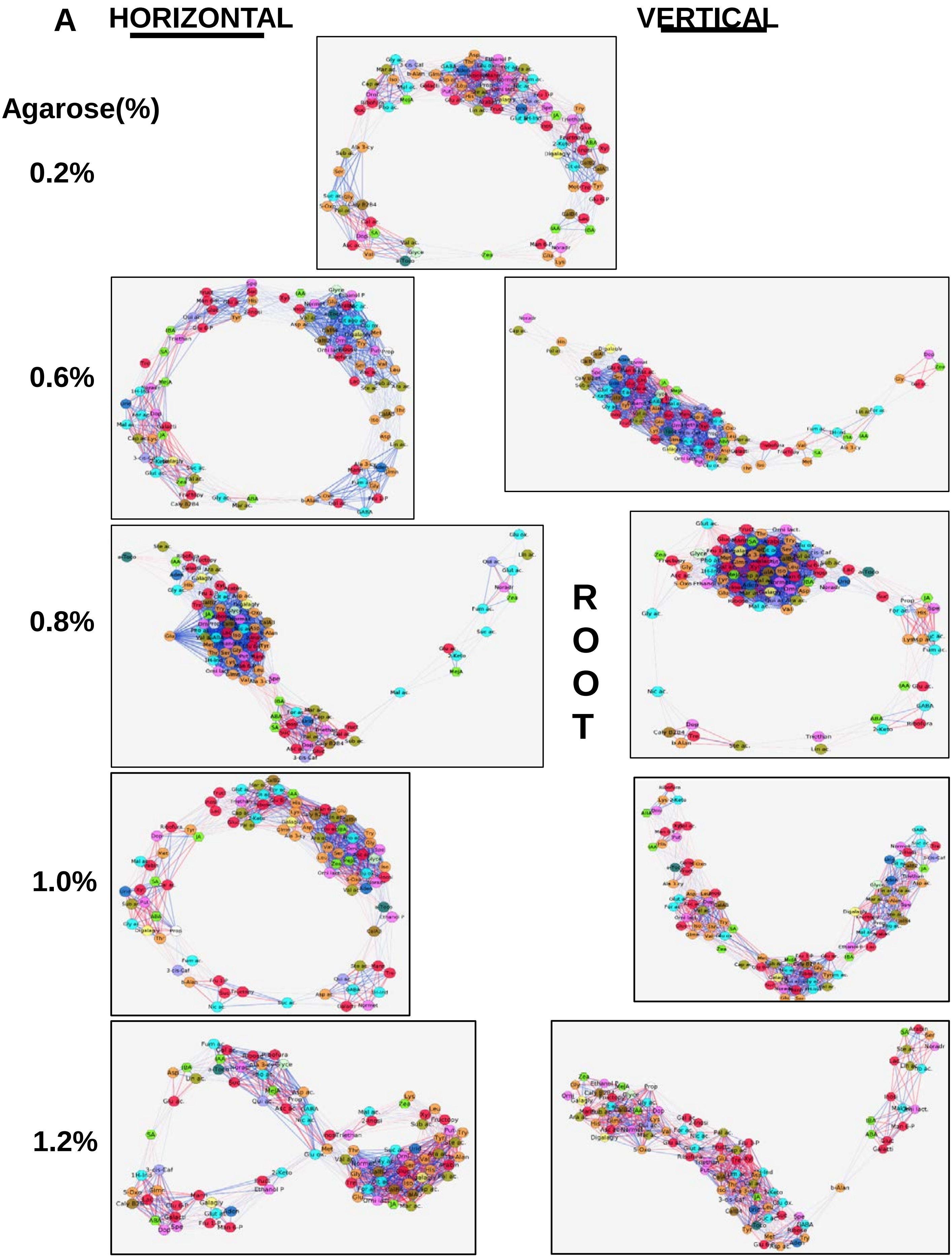

Supplementary Figure S5

**B** **HORIZONTAL** **VERTICAL**

**Agarose(%)**

**0.2%**

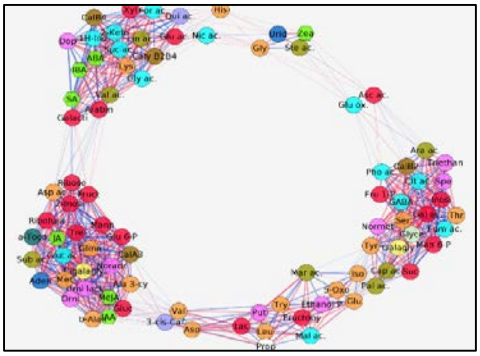

**0.6%**

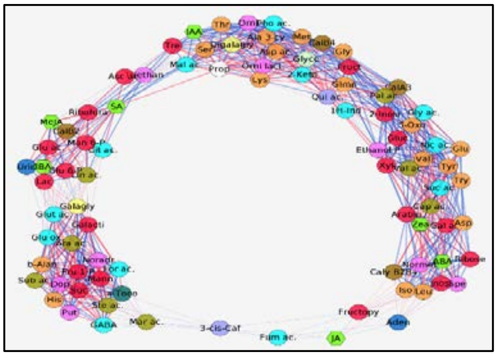

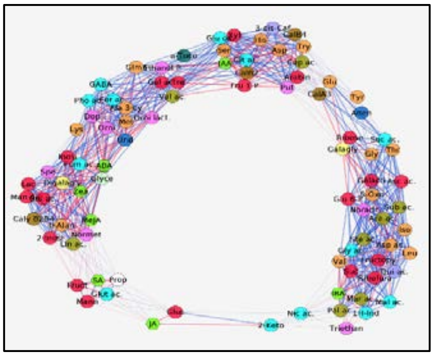

**SHOOT**

**0.8%**

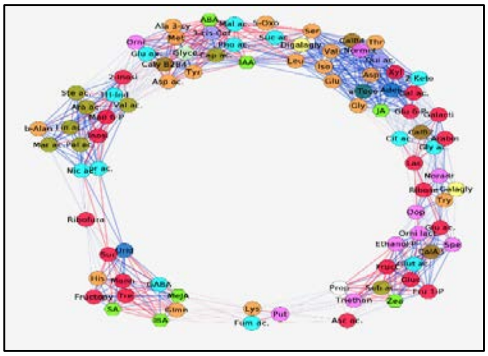

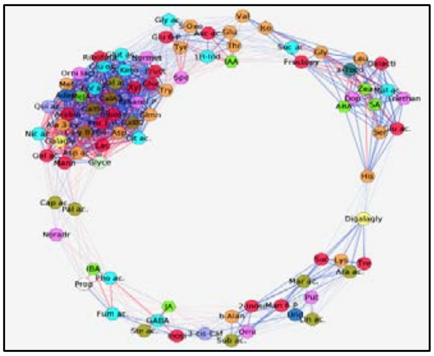

**1.0%**

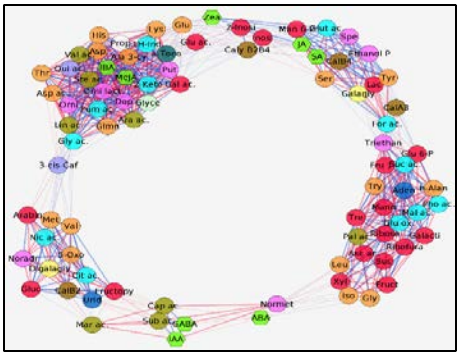

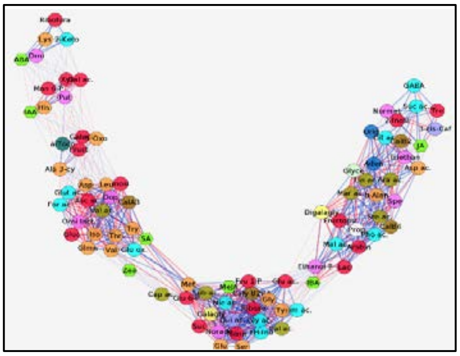

**1.2%**

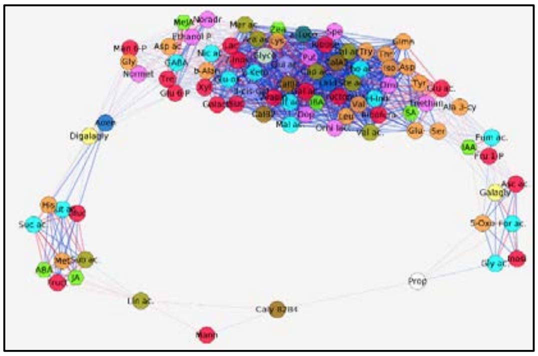

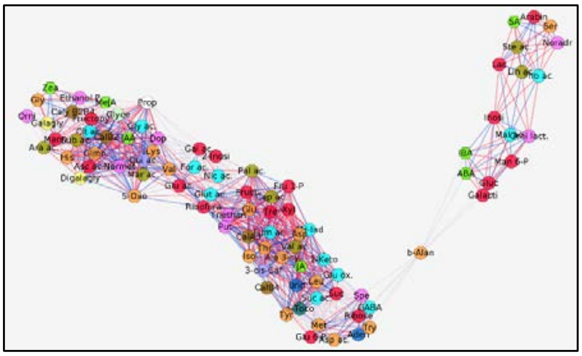

### Supplementary Figure S5

**Figure S5: Impact of increasing hardness on metabolite/hormone correlation networks.**

**A,** Root. **B,** Shoot. Networks were drawn using Cytoscape software. The blue and red lines between the different nodes indicate positive and negative correlations respectively. Metabolites not mapping in any cluster are connected with faint lines. In networks, sugars (red), amino acids (golden yellow), organic acids (cyan), lipids (light yellow), fatty acids (light brown), monoamines (pink), nucleotides (light blue), hydroxycinnamic acids (light purple), alkane (ash colour) and phytohormones (fluorescent green hexagon) are represented. Interactions with $r \geq \pm 0.95$ were used for generating network

### Supplementary Figure S6

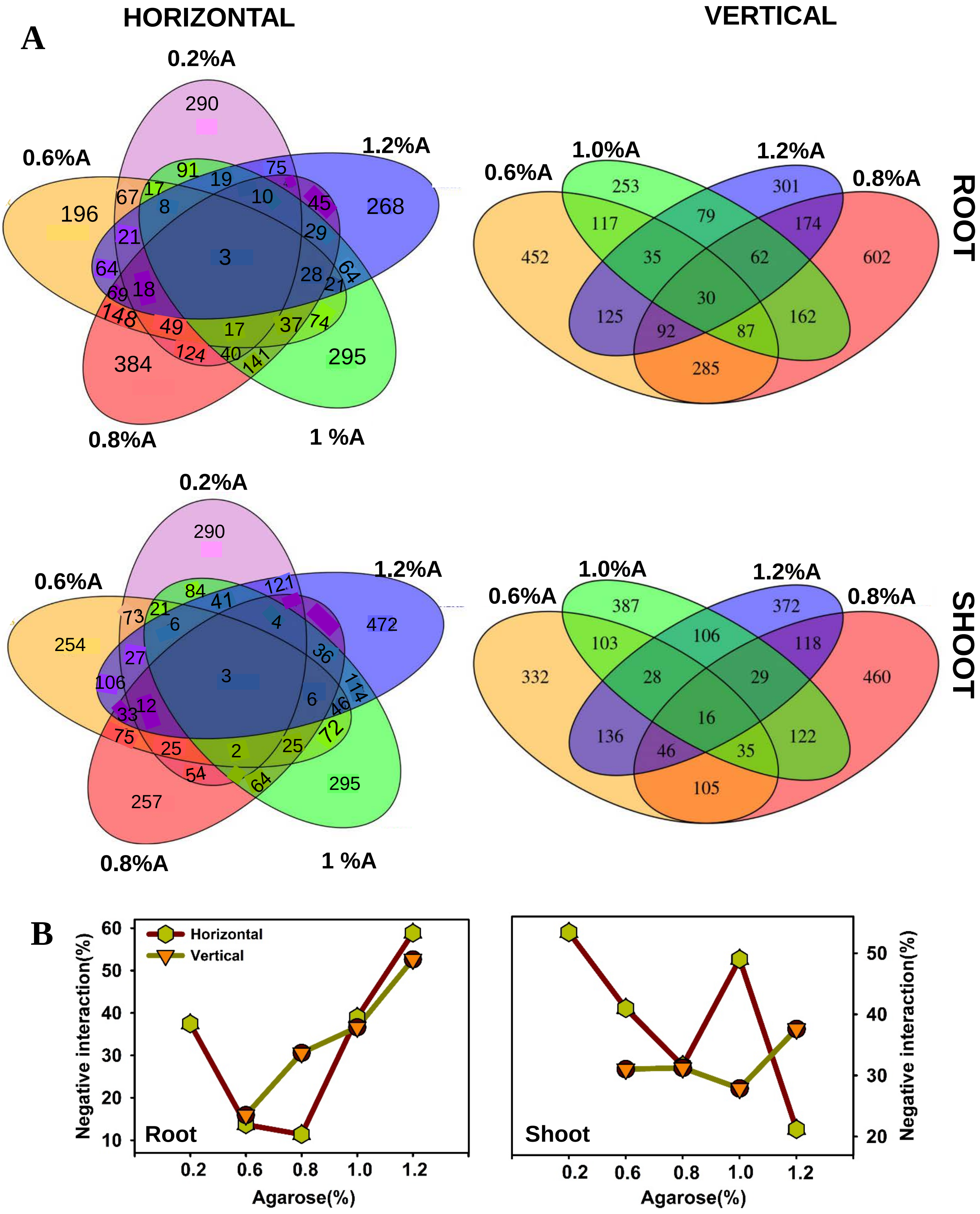


**Figure S6: Reduction in shared interaction among metabolites with increasing hardness.** **A.** Venn diagrams depicting unique and shared interactions. **B,** Increase in negative interactions in roots with increasing hardness. **Note-** Compared to vertically-impeded shoot/root, the horizontally impeded root/shoot in Venn diagram shows very few shared interaction at all hardness. Venn diagrams illustrating the number of shared and unique interactions in the metabolic networks were prepared using Pearson's Correlation coefficient (PCC) value ± ≤ 0.95.
