## Supplementary material for "Tomato roots sense horizontal/vertical mechanical impedance and divergently modulate root/shoot metabolome": Table S1

**Table S1: List of genes and the primers used for qRT-PCR analysis.** The primers were designed using SGN SL2.50 version (<https://solgenomics.net>). MCA, β-actin and ubiquitin were from NCBI Unigene.

| **Name of gene/pathway** | **Gene/Sol I.D.** | **Primer Sequence (5ʹ🡪3ʹ)** | | | **Start position** | **End position** | **Amplicon size (bp)** |
| --- | --- | --- | --- | --- | --- | --- | --- |
| *TOR* | Solyc01g106770.2 | Fp | CCAACTTGTCTCAAAGGAGCTTA | | 86846987 | 86847009 | 102 |
|  |  | Rp | TTGTGTTCACCAAAATACAGACG | | 86847066 | 86847088 |  |
| *RAPTOR* | Solyc09g014780.2 | Fp | GCCAAATCTGTTAAGTCAACTGG | | 6952512 | 6952534 | 115 |
|  |  | Rp | GTCCTCCATTCATATCAAACCAA | | 6952589 | 6952609 |  |
| *Lst8* | Solyc03g059310.2 | Fp | AGTCAAGATTTGGAATGTGGATG | | 24658355 | 24658378 | 120 |
|  |  | Rp | TCAGAAGATGCTGTGATGAGAAA | | 24658571 | 24662207 |  |
| *S6k* | [Solyc03g095510.2.1](https://solgenomics.net/feature/17765077/details) | Fp | CAAAGAGTTGGATCTCGTGTTCT | | 51201291 | 51201313 | 116 |
|  |  | Rp | GCATAATGCACAGGGATCTAAAG | | 51201588 | 51201610 |  |
| Hexokinase 1 (*HXK1*) | Solyc03g121070 | Fp | CCTCCTTGAAGACGAGAAAACT | | 63740763 | 63740784 | 117 |
|  |  | Rp | GCGTATCTCTTCCCATCTTTTT | | 63740858 | 63740879 |  |
| Sucrose synthase  (*SUS3*) | Solyc07g042550 | Fp | TCTTGATGCTACTTTGGATGCT | | 55824505 | 55824526 | 100 |
|  |  | Rp | CAGTAGCTGGTGAGGTTTCAAG | | 55825021 | 55825042 |  |
| *Invertase* (*Lin6*) | Solyc10g083290 | Fp | CCGCTATCTACCCGTCTAAAGT | | 62274577 | 62274598 | 113 |
|  |  | Rp | GTTAGCATCCACAATTCCAGTG | | 62274668 | 62274689 |  |
| *Invertase 8* | Solyc10g083300 | Fp | GGGCTCTTAACTTTGGCTTCTA | | 62286241 | 62286262 | 103 |
|  |  | Rp | CATCAGAGCACATGAGAACCTT | | 62286322 | 62286343 |  |
| *MYB36* | Solyc07g006750 | Fp | AGTTAACCCCTCTCCAACAAAA | | 1562031 | 1562052 | 107 |
|  |  | Rp | AGCATGGTTTTCACTAGGGTTT | | 1562116 | 1562137 |  |
| Hy5 (*Hy5*) | Solyc08g061130 | Fp | CTAGTTTCGGGTGGATTGTTCT | | 44862061 | 44862082 | 104 |
|  |  | Rp | GCTTAGCCACAATCCTTCACTC | | 44862143 | 44862164 |  |
| (MCA1) *(*mid1-complementing activity 1) | Unigene id SGN-U565830 | Fp | GCCCTCCTGCTGATCTGTGC | | 44901207 | 44901227 | 153 |
|  |  | Rp | CCCAATTAGCCTCACCGCATC | | 44901341 | 44901360 |  |
| Sucrose transporter *(SUS* *trans*) | Solyc11g017010 | Fp | ACTCGGTATTCCTCATCGATTT | | 7859205 | 7859226 | 112 |
|  |  | Rp | GAAACGTGAGGAGCAATTATCA | | 7859295 | 7859316 |  |
| SnRK1-interacting protein 1 (*SNRK1*) | Solyc11g040110 | Fp | | GAGATCCTACGTGACCTGAACA | 38626936 | 38626957 | 116 |
|  |  | Rp | | TAGAAGCTCAACATCCGGTTAG | 38627269 | 38627290 |  |
| Regulator of G protein signaling (*RGS1*) | Solyc05g014160 | Fp | | ACCACTCTTCCGAAGCTCTTAT | 7970615 | 7970636 | 103 |
|  |  | Rp | | TACTGCCTTTGGACCTAGGAAT | 7969976 | 7969997 |  |
| E2F (*E2F*) | Solyc01g007760 | Fp | | TAGCAGTTAAAGCTCCTCATGG | 1871489 | 1871510 | 108 |
|  |  | Rp | | GACCCATTGTGCTTCTGAGTAT | 1871651 | 1871672 |  |
| Trehalose-6-Phosphate  (*Tre1*) | Solyc03g083960 | Fp | | GTAGTCACGGCATGGATATAA | 48481577 | 48481598 | 117 |
|  |  | Rp | | CTCACTAGCAGGTTGGAAAAGA | 48481768 | 48481788 |  |
| Trehalose -6-phosphate  (*Tre*2) | Solyc06g060600 | Fp | | GCGATGACAGAACAGATGAAGA | 36273615 | 36273636 | 105 |
|  |  | Rp | | AGTATCCCCGAGAGAAAAGACA | 36273795 | 36273816 |  |
| β-ACTIN | FJ532351.1 Unigene | Fp | | GTCCCTATTTACGAGGGTTATGC | 278213 | 278235 | 72 |
|  |  | Rp | | CAGTTAAATCACGACCAGCAAGATT | 278163 | 278186 |  |
| UBIQUITIN 1 | 3/X58253. Unigene | Fp | | GCCGACTACAACATCCAGAAGG | 69850003 | 69850024 | 142 |
|  |  | Rp | | TGCAACACAGCGAGCTTAACC | 69849882 | 69849902 |  |
